# Single-Plant Genome-Wide Association Study Identifies Loci Controlling Multiple Vegetative Architecture Traits in Cultivated Northern Wild Rice (*Zizania palustris L*.)

**DOI:** 10.64898/2026.04.15.718548

**Authors:** L McGilp, R. Millas, A. Mickelson, L.M. Shannon, J. Kimball

**Affiliations:** Department of Agronomy and Plant Genetics, University of Minnesota, St. Paul, Minnesota, USA; Department of Horticultural Science, University of Minnesota, St. Paul, Minnesota, USA

**Keywords:** plant architecture, polygenic regulation, comparative genomics, single nucleotide polymorphism

## Abstract

Cultivated Northern Wild Rice (*Zizania palustris* L.) is an obligately outcrossing, self-incompatible cereal grown in aquatic paddies in the United States. Genetic improvement has relied primarily on phenotypic recurrent selection, and genomic approaches remain largely unexplored in this emerging crop. We applied a single-plant genome-wide association study (sp-GWAS) framework to dissect vegetative architecture traits in five open-pollinated cultivated populations evaluated across three years (n = 2,173 plants). Plant height (PH), basal stem width (BSW), primary stem width (PSW), flag leaf length (FLL), and flag leaf width (FLW) were analyzed using a mixed linear model accounting for population structure and kinship. Broad-sense heritability ranged from 0.03 to 0.34, and year effects explained up to 54% of phenotypic variance, indicating strong environmental influence. After filtering 73,363 SNPs, genome-wide linkage disequilibrium decayed rapidly (r² = 0.1 at ∼2.3 kb). A total of 124 significant SNPs (FDR < 0.01) were consolidated into 98 loci, of which 46 were associated with multiple traits and 11 were shared across four traits. Candidate genes near multi-trait loci included conserved regulatory classes implicated in grass architecture, including *HLH/bHLH* transcription factors. Diplotype analyses at candidate loci revealed both simple biallelic and complex multi-allelic haplotype structures, indicating that locus-level haplotype effects underlie several GWAS signals. Results demonstrate that sp-GWAS can detect statistically robust associations in a highly heterozygous, non-replicable crop system and suggest a polygenic, coordinated genetic architecture governing vegetative growth. These findings support genomic prediction and multi-trait selection strategies to accelerate improvement of cultivated Northern Wild Rice.

**PLAIN LANGUAGE SUMMARY:** Cultivated Northern Wild Rice is an important specialty crop grown in flooded paddies in the United States. Unlike many major crops, it is naturally outcrossing and highly variable, which makes traditional breeding challenging and slow. Most improvement efforts have relied on selecting plants based only on how they look in the field, and genomic tools have rarely been used. In this study, we used DNA markers to better understand the genetics behind plant structure traits such as plant height, stem thickness, and leaf width. We evaluated more than 2,000 plants from five cultivated populations over three growing seasons. Because weather and growing conditions strongly influence these traits, we used statistical models to separate environmental effects from genetic effects. We identified 98 regions of the genome associated with variation in plant structure. Many of these regions influenced more than one trait, showing that plant height, stem strength, and leaf size are genetically connected. Several regions contained genes similar to those known to control plant growth and development in other grasses. We also found that, in some cases, combinations of nearby DNA variants (haplotypes) explained trait differences better than single genetic markers. Overall, this work shows that modern genomic tools can successfully identify useful genetic variation in cultivated Northern Wild Rice, even though it is highly outcrossing and genetically diverse. These results provide a foundation for using genomic selection to improve plant structure, lodging resistance, and overall performance in breeding programs.

**CORE IDEAS:**

- Single-plant GWAS successfully detects genetic associations in obligately outcrossing cultivated Northern Wild Rice where conventional replicated mapping populations are impractical.
- Vegetative architecture traits exhibit low heritability but retain recoverable polygenic signal, where nearly half of detected loci influence multiple architecture traits, indicating integrated developmental control.
- Genome-wide linkage disequilibrium decays rapidly (∼2.3 kb), consistent with expectations for an obligately outcrossing species and supporting relatively localized association signals.
- Candidate genes include conserved regulatory classes (*TE1*-like, *HLH/bHLH*, *SPL*).
- Given extensive overlap between QTL and environmental effect, multi-trait, multi-environment genomic prediction provides a pragmatic breeding strategy to improve canopy efficiency, lodging resistance, and harvestability in aquatic production systems.

## INTRODUCTION

Cultivated Northern Wild Rice (cNWR; *Zizania palustris* L.) is a high-value, small commodity crop grown in irrigated paddies, primarily in Minnesota and California in the United States. The ongoing domestication of cNWR and development of varieties with economically desirable phenotypes have focused on yield, plant vigor, resistance to seed shattering and diseases, uniformity characteristics and seed size (Grombacher et al., 1997; McGilp et al., 2023; Porter, 2019). Breeders of cNWR utilize phenotypic-based recurrent selection for population improvement, which is time-consuming, limits the selection opportunities compared to other advanced breeding schemes, and does not take advantage of the numerous genomic technologies available to researchers. Therefore, it is essential to move towards more efficient breeding methods which can expedite cultivar development.

An understanding of the genetic architecture which underlies agronomic traits, through biparental mapping and Genome-wide Association Studies (GWAS), has facilitated crop improvement in many species (Ain et al., 2015; Alqudah et al., 2014; Atwell et al., 2010; X. Huang et al., 2010, 2012; M. Wang et al., 2012). However, only two biparental mapping studies have been undertaken in NWR (Kahler et al., 2014; Kennard et al., 2002). This limited progress reflects the challenges associated with developing and maintaining biparental mapping populations. These challenges stem from biological constraints, including unorthodox seed storage requirements that limit viability to under two years in storage (Grombacher et al., 1997; Oelke & Porter, 2016) and self-incompatibility mechanisms that reduce seed set and plant viability over generations of selfing.

An alternative to biparental mapping is GWAS, which leverages historical recombination and standing genetic variation to identify loci associated with complex quantitative traits (Huang et al., 2009; Huang & Han, 2014). The high levels of genetic diversity present within cultivated and wild populations of NWR make GWAS a particularly attractive approach for trait discovery (McGilp et al., 2025). Recent advancements in *Z. palustris* genomic resources have made GWAS studies in the species more feasible. These include an annotated chromosome-scale reference genome (Haas et al., 2021) and genome-wide SNP markers generated through genotyping-by-sequencing (GBS) to characterize population structure and genetic diversity across wild and cultivated NWR in the Great Lakes region (McGilp et al., 2025; Shao et al., 2019). Together, these resources establish the necessary foundation for GWAS in NWR and create new opportunities to link phenotypic variation with underlying genetic mechanisms.

Traditional GWAS approaches in self-pollinating species often utilize clonally replicated or highly inbred lines to reduce within-genotype variability and improve statistical power (Atwell et al., 2010; Brachi et al., 2010; Huang et al., 2010; Wang et al., 2012). However, GWAS has also been successfully applied in outcrossing species and natural populations where genetically identical replicates are not feasible (Hayward et al., 2016; Karczewski et al., 2024; Klein et al., 2005; Marina et al., 2021; McCarthy et al., 2008; Omeka et al., 2022; Pairo-Castineira et al., 2023). To address this challenge explicitly in crops, a single-plant GWAS (sp-GWAS) framework was recently developed and validated in maize (*Zea mays*) (Gyawali et al., 2019) and luffa (*Luffa* spp.) (Y.-D. Li et al., 2024). In sp-GWAS, phenotypes are collected from individual plants without replication, and statistical power is recovered through large sample sizes and mixed models that account for kinship, population structure, and environmental variation. For NWR, sp-GWAS presents a practical and scalable strategy for genetic mapping that aligns with the biological realities of an obligately outcrossing species.

In this study, we apply a sp-GWAS framework to investigate the genetic architecture of vegetative morphology in cNWR. We focus on plant height, basal stem width, primary stem width, flag leaf length, and flag leaf width, which are traits that can be readily collected on a large population of individuals, are widely studied in other cereal crops, and are directly relevant to cNWR breeding goals. Collectively, these traits influence plant vigor, canopy structure, lodging resistance, and harvestability in aquatic production systems, where excessive height or insufficient stem strength can reduce yield stability. By investigating the genetic associations underlying vegetative architecture, this work aims to evaluate the utility of sp-GWAS in cNWR and to provide a foundation for genomics-enabled improvement of this emerging crop.

## MATERIALS AND METHODS

### Plant Materials

Five open-pollinated populations of cNWR were utilized in this study including: ‘K2’ (released in 1972), the third released cNWR variety, ‘Barron’ (released in 2014), an early-maturing variety with improved resistance to fungal brown spot (Bipolaris oryzeae); ‘FY-C20’, an elite breeding population selected for long seed length; and two cultivars, ‘Itasca-C12’ and ‘Itasca-C20’ (released in 2007 and 2019, respectively), developed through recurrent selection of the ‘Itasca’ cultivar (released in 2000). Varieties derived from the Itasca pedigree have been the industry standard in cNWR for the past 25 years.

All populations were grown from 2019 to 2021 at the University of Minnesota’s North Central Research and Outreach Center experimental paddies in Grand Rapids, MN, USA (47.2425°N, 93.4927°W; average annual temperature 4.64 °C; annual precipitation 73.61 cm; elevation 392Lm). In May of each year, seeds were sown in large plots (24L×L12Lm), each consisting of 36 24Lm-long rows spaced 0.38Lm apart. Sowing depth was approximately 2.54 cm, at a rate of ∼1 plant per ft² (∼18 plants per m²), totaling ∼3,200 plants per population. Paddies were flooded to an average depth of 20Lcm after sowing. Prior to the aerial growth stage of NWR (Duquette & Kimball, 2020), copper sulfate (Chem One, Houston, TX; 1.13LkgLhaL¹) was applied to manage algal growth, and Aquabac (*Bacillus thuringiensis* var. Israelensis, Becker Microbial Products, Coral Springs, FL; 1.86LkgLhaL¹) was used to control midge (Diptera) and leaf miner. Fungicides and insecticides were applied during the reproductive stage to manage *Bipolaris* spp. and riceworm (*Apamea apamiformis*). Water was slowly drained in early to mid-August to facilitate harvest. Temperature and precipitation data were collected in Grand Rapids, MN between May and September of each year in this study and monthly averages of min, max, and average daily temperature and precipitation were calculated.

Two hundred plants from each population were randomly selected and tagged to form the association panel. Due to the high heterozygosity of cNWR germplasm, genotypically identical plants could not be assessed across years. At flowering, the leaves below the flag leaves were collected from each plant and stored at −80L until transport to the UMN St. Paul Campus for DNA extraction and genotyping.

### Phenotyping

For each of the three years, five morphological traits were measured for each plant in the association panel, including: plant height (PH), the length in cm from the soil line to the tip of the female panicle; basal stem width (BSW), the width in mm of the base of the primary stem; primary stem width (PSW), the width in mm of the primary stem at the flag leaf; flag leaf length (FLL), the length in cm from the base of the flag leaf to its tip; and flag leaf width (FLW), the width in cm of the flag leaf at its widest point (Supplemental Table S1). The number of plants phenotyped for each trait, population, and year varied when accounting for plant mortality or missing data (Supplemental Table S2).

Because cNWR is obligately outcrossing and individual genotypes cannot be clonally propagated or re-evaluated across years, genotypically identical plants were not observed in multiple environments. For statistical modeling, year and population were therefore modeled as random effects to account for environmental and background genetic variance, while allowing inference beyond the specific years and populations sampled. Variance components were estimated using the lme4 R package version 1.1-35.5 (Bates et al., 2015). Means and standard errors were calculated both across years and within years. Trait correlations were calculated using Hmisc version 5.2.3 (Harrell, 2025). Broad-sense heritability was calculated using the equation 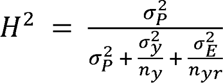 as described in (Holland et al., 2002) where 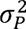 = genetic variance between populations, 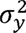 = variance due to year, 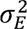= residual variance, n_y_= harmonic mean of the number of years in which each population is observed and n_yr_= harmonic mean of the number of replications across years.

### Genotyping-By-Sequencing and Variant Calling

From each leaf sample, ∼20 mg of tissue was freeze-dried and ground using a TissueLyser II (Qiagen, Valencia, CA, USA) at 25 Hz for 5 minutes. Genomic DNA was extracted using the DNeasy Plant Mini Kit (Qiagen, Valencia, CA, USA), following the manufacturer’s instructions. DNA concentration and quality was assessed with a NanoDrop spectrophotometer (Thermo Scientific, Wilmington, DE, USA). DNA libraries for genotyping-by-sequencing (GBS) were prepared at the University of Minnesota Genomics Center (UMGC; Saint Paul, MN, USA; http://genomics.umn.edu). Library preparation followed the protocol of (Elshire et al., 2011). Genomic DNA was digested using *Pst*I and *Msp*I and ligated to barcoded adapters. Paired-end sequencing (2 x 150 bp) was performed on a single lane of an Illumina NovaSeq 6000 flow cell, targeting 2.25B reads per lane (∼2M reads per sample).

Sequence data for each year were processed separately. Adapter trimming was performed using Cutadapt version 4.6 (Martin, 2011), data quality was assessed with FastQC version 0.11.7 (Andrews, 2010) and MultiQC version 1.9 (Ewels et al., 2016). Reads were aligned to the *Z. palustris* reference genome v1.0 (Haas et al., 2021) using Burrows-Wheeler Aligner Maximal Exact Match Aligner (BWA-MEM) version 0.7.17 (Li, 2013). SNP calling was performed with BCFtools version 1.20 (Danecek et al., 2021) with a minimum mapping quality of 20 through GNU parallel (Tange, 2021). SNPs were filtered with BCFtools for a minor allele frequency (MAF) ≥ 0.03, a minimum depth of 8, no indels, only biallelic SNPs and a maximum of 10% missing data. VCF files were subsetted to include only the top 17 scaffolds corresponding to the 15 chromosomes and two unplaced scaffolds based on Haas et al. (2021). Low quality samples as well as samples without phenotypic data were removed. Genotype imputation was performed using Beagle v5.4 (Browning et al., 2021). Following imputation, BCFtools was used to make a VCF file containing only SNPs that were present across all three years, which was used for all future steps. Tassel 5 (Bradbury et al., 2007) was used to generate a HapMap file. Unless otherwise stated, all programs and packages were used according to their default parameters.

### Population Structure and Linkage Disequilibrium

A total of 73,363 filtered SNP markers were used for analysis of the genotypic data. Principal component (PC) analysis (PCA) was conducted in PLINK version 1.90 (Chang et al., 2015) and visualized with ggplot2 version 3.5.1 (Wickham, 2016) R package. Linkage disequilibrium (LD) decay was calculated using PopLDdecay version 3.43 (C. Zhang et al., 2019) on a per chromosome and genome-wide basis for the full SNP data set as well as a data set filtered for genic regions. We report genome-wide and chromosome-level LD decay at r² = 0.1.

### Association Analysis

Genome-wide association analyses were performed using GAPIT (Wang & Zhang, 2021), with a Mixed Linear Model (MLM). The model was: y = µ + Q + K + e, where *y* is the phenotype vector, μ is the overall mean, Q is the population structure matrix (first 4 PCs), *K* is the kinship matrix (VanRaden method) (VanRaden, 2008), and *e* is the residuals. A false discovery rate (FDR) threshold of < 0.01 was used to determine significance. Significant SNPs were clustered into loci using chromosome-specific LD decay distances (significant SNP ± decay window).

### Candidate Gene and Diplotype Analysis

Candidate genomic regions underlying significant GWAS loci were further examined using a diplotype-based approach. For each locus of interest, SNPs located within ±3 kb of the boundaries of the significant SNPs were extracted from the full variant dataset. Genotype data were phased using Beagle to infer haplotype structure within each region. Phased SNP data were then subset into locus-specific VCF files for downstream analysis. Diplotypes were defined based on the allelic state across all SNPs within each locus-specific window and were retained for analysis if represented by more than 25 individuals. Diplotype classes were manually curated and visualized to assess local haplotype structure. Phenotypic distributions of each diplotype were summarized using boxplots. Statistical differences among diplotypes were evaluated separately for each trait using ANOVA. When significant effects were detected, mean separation was performed using Tukey’s honestly significant difference (HSD) test as implemented in the agrocolae package v1.3-7 (de Mendiburu, 2023). All data processing, visualization, and statistical analyses were conducted in R. Figures were generated using ggplot2 and ggpubr v0.60 (Kassambara, 2023), with additional custom scripts developed to visualize diplotype structure and phenotypic distributions.

To further characterize genes within each significant locus, predicted protein sequences from annotated gene models were extracted and subjected to BLASTP searches against the NCBI non-redundant (nr) protein database. Protein similarity searches were conducted using BLAST+ with default parameters, and top hits were retained based on E-value (<1e−5), sequence identity and alignment coverage. Conserved domains were identified using NCBI’s Conserved Domain Database (CDD) to support functional inference. Putative gene functions were integrated with GWAS and diplotype results to prioritize biologically plausible candidate genes underlying trait variation.

## RESULTS

### Weather and Climate

From 2019 (Year 1) to 2021 (Year 3), temperature and precipitation patterns from May through September showed consistent northern temperate seasonal trends with some interannual variability. May was consistently the coolest month, with average temperatures between 9.44°C in Year 1 and 12.48°C in Year 3, while July was the warmest, with average temperatures ranging from 20.38°C in Year 1 to 21.49°C in Year 3 (Supplemental Table S3). Average precipitation levels were generally low across all years (< 0.3 inches), with May being the driest month (Supplemental Table S3). August 2020 (Year 2) had the highest recorded precipitation (0.28 inches), whereas the summer of Year 3 was notably dry, with minimal rainfall in June (0.04 inches) and August (0.06 inches). Overall, Year 3 was the warmest and driest year, while Year 1 showed slightly cooler temperatures and higher early-season precipitation.

### Phenotypic Variation Among Cultivated Populations

Five open-pollinated populations of cNWR were evaluated for plant height (PH), basal stem width (BSW), primary stem width (PSW), flag leaf length (FLL), and flag leaf width (FLW) across three years, totaling 2,173 phenotyped individuals (∼145 plants per population per year) (Supplemental Table S2). Year and population were modeled as random effects. Year explained the largest proportion of variance for all traits (6.8 - 54.1%), while population effects were small (0.36 - 2.95%), indicating strong environmental influence but detectable genetic differentiation (Table 1). When averaged across years, PH ranged from 172.1 cm in Itasca-C12 to 189.6 cm in K2; FLL ranged from 35.4 cm in Barron to 38.5 cm in Itasca-C20; FLW was lowest in Itasca-C12 (2.3 cm) and highest in both Itasca-C20 and K2 (2.5 cm); PSW was lowest in FY-C20 and Itasca-C12 (6.3 mm) and highest in K2 (6.5 mm); and BSW ranged from 12.8 mm in FY-C20 to 13.5 mm in Itasca-C20 (Table 1). Substantial phenotypic variation was observed between years, with the lowest trait values consistently occurring in Year 2 and the highest in Year 3. Across genotypes, yearly trait means ranged from 152.5 to 204.0 cm for PH, 10.8 mm - 14.6 mm for BSW, 5.6 - 7.3 mm for PSW, 34.3 - 39.2 cm for FLL, and 2.1 - 2.7 cm for FLW (Supplemental Table S4).

**Table 1.** Proportion of phenotypic variance explained by year and population, and mean trait values (± SE) of vegetative architecture traits across years and populations of cultivated Northern Wild Rice (cNWR; *Zizania palustris* L.); variance components were estimated using a linear mixed-effects model with year (YR) and population as random effects, and values represent the percentage of total phenotypic variance attributable to year, population, and residual error. Year means (YR1–YR3) are averaged across populations, and population means (Barron, FY-C20, Itasca-C12, Itasca-C20, and K2) are averaged across years. Plant height (PH), flag leaf length (FLL), and flag leaf width (FLW) are reported in centimeters (cm), and primary stem width (PSW) and basal stem width (BSW) are reported in millimeters (mm).

|  | % Variation Explained |  |  | Variation by Year (YR) |  |  | Variation by Population |  |  |  |  |
| --- | --- | --- | --- | --- | --- | --- | --- | --- | --- | --- | --- |
| Trait | Year | Population | Residual | YR1 | YR2 | YR3 | Barron | FY-C20 | Itasca-C12 | Itasca-C20 | K2 |
| PH | 54.13 | 2.95 | 42.92 | 191.00 ± 2.00 | 152.46 ± 1.49 | 204.02 ± 1.41 | 188.15 ± 1.81 | 183.17 ± 1.74 | 172.05 ± 1.29 | 179.88 ± 1.28 | 189.55 ± 0.92 |
| FLL | 6.75 | 1.26 | 91.99 | 34.59 ± 0.87 | 34.33 ± 0.71 | 39.19 ± 0.73 | 35.40 ± 0.46 | 35.57 ± 0.47 | 36.23 ± 0.47 | 38.45 ± 0.48 | 35.76 ± 0.41 |
| FLW | 20.93 | 1.31 | 77.76 | 2.41 ± 0.04 | 2.12 ± 0.04 | 2.72 ± 0.04 | 2.48 ± 0.03 | 2.38 ± 0.03 | 2.32 ± 0.02 | 2.49 ± 0.03 | 2.49 ± 0.02 |
| PSW | 32.97 | 0.36 | 66.68 | 6.31 ± 0.10 | 5.63 ± 0.07 | 7.26 ± 0.08 | 6.50 ± 0.06 | 6.33 ± 0.06 | 6.33 ± 0.05 | 6.45 ± 0.06 | 6.53 ± 0.05 |
| BSW | 37.37 | 0.56 | 62.08 | 14.12 ± 0.22 | 10.80 ± 0.15 | 14.57 ± 0.20 | 13.01 ± 0.15 | 12.82 ± 0.14 | 13.35 ± 0.14 | 13.45 ± 0.14 | 13.12 ± 0.11 |

All pairwise trait correlations were positive and statistically significant (p-value ≤ 0.05) (Supplemental Table S5), indicating coordinated variation in vegetative architecture. Stem and leaf width traits were particularly strongly correlated, with PSW-FLW (r = 0.75), PSW-BSW (r = 0.60), and FLW-BSW (r = 0.51) showing tight relationships. Plant height was moderately correlated with both stem widths (PH-PSW r = 0.55; PH-BSW r = 0.59) and leaf width (PH-FLW r = 0.50), suggesting shared growth regulation. In contrast, flag leaf length (FLL) exhibited weaker correlations with other traits (e.g., PH-FLL r = 0.22).

Broad-sense heritability was estimated for each trait using harmonic means to account for the unbalanced dataset (Table 2). Heritability estimates were low for most traits including 0.14 for PH, 0.04 for BSW, 0.03 for PSW, 0.34 for FLL, and 0.16 for FLW. Only FLL showed moderate heritability, suggesting greater genetic control, while the remaining traits showed low heritability estimates, reflecting substantial environmental variance or limited detectable genetic variance within this panel under single-plant sampling.

**Table 2.** Variance components and broad-sense heritability (H²) estimates for vegetative architecture traits in cultivated Northern Wild Rice (*Zizania palustris* L.); variance components were estimated using a linear mixed-effects model with year and population as random effects. Genetic variance represents variance attributable to population, environmental variance includes year and residual components, and phenotypic variance is the total observed variance. Broad-sense heritability (H²) was calculated using harmonic means to account for unbalanced sampling across years and populations. Units are indicated for each trait.

| Trait | Genetic Variance | Phenotypic Variance | Environmental Variance | Broad-sense heritability ($H^2$) |
| --- | --- | --- | --- | --- |
| Plant height (cm) | 39.07 | 717.79 | 569.19 | 0.14 |
| Basal stem width (mm) | 0.06 | 4.16 | 6.91 | 0.04 |
| Primary stem width (mm) | 0.01 | 0.66 | 1.34 | 0.03 |
| Flag leaf length (cm) | 1.37 | 7.37 | 100.45 | 0.34 |
| Flag leaf width (cm) | 0.01 | 0.09 | 0.32 | 0.16 |

### Genotypic Data Structure and Linkage Disequilibrium

A total of 73,363 high-quality, filtered SNPs shared across all three years of genotyping for 2,173 samples were retained for downstream analysis. Average marker spacing was 16.7 kb genome-wide, with chrs 1, 2, 6, and 13 harboring the highest SNP densities (Table 3). LD decay was assessed using the full set of SNPs (Figure 1A) as well as a subset restricted to genic regions (Figure 1B). At an *r^2^* threshold of 0.1, the genome-wide LD decay distance was 2,300 bp for the full dataset and 3,500 bp for the genic subset. Chromosome-specific LD decay distances for the full dataset ranged from 800 to 2,800 bp (mean = 1,747 bp), while for the genetic subset, LD decay ranged from 70 bp to 12,500 bp (mean = 2,310 bp) (Table 3).

**Figure 1.**
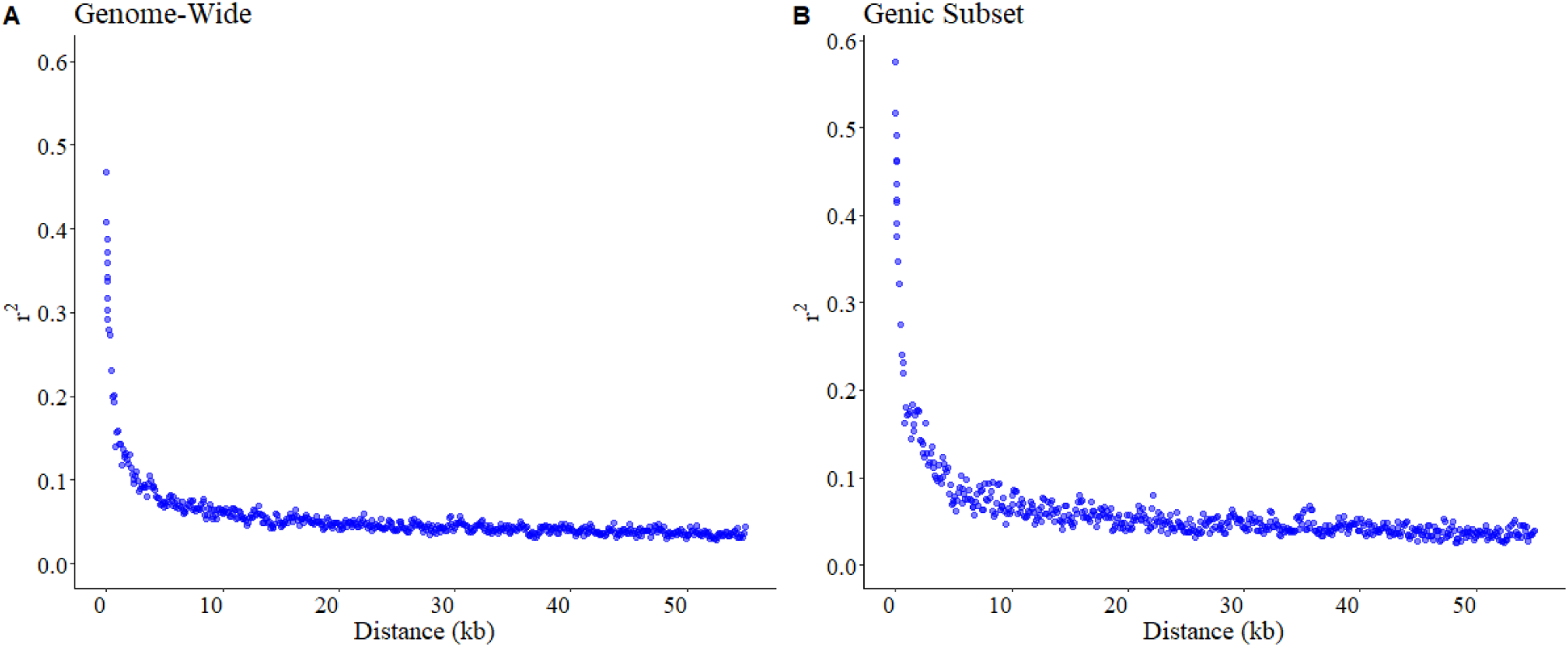
Linkage disequilibrium (LD) decay in cultivated Northern Wild Rice (cNWR; *Zizania palustris*) based on (A) all filtered genome-wide SNPs (n = 73,363) and (B) SNPs located within annotated gene models (genic subset). Pairwise LD (r²) is plotted against physical distance (kb). Points represent mean pairwise r² values within distance bins. PCA was conducted separately for each individual year and on the final merged dataset containing overlapping SNPs across years (Figure 2). In all cases, PC1 clearly separated K2 from the other cNWR populations, explaining 27.7% (Year 2) to 35.3% (combined years) of the total genotypic variation. The remaining populations formed a continuous cluster along PC1, with Barron and both Itasca populations distinguishable from one another. The second PC explained 11.7% (Year 2) to 17.6% (Year 1) of the total variation and further separated Barron from the Itasca populations, with FY-C20 positioned intermediately. Further examination of the final merged dataset using additional principal components (Supplemental Figure S1) revealed finer-scale structure. In PC2 vs. PC3, FY-C20 and Barron formed distinct clusters, and all comparisons involving PC1 consistently isolated K2 as a separate cluster.

**Figure 2.**
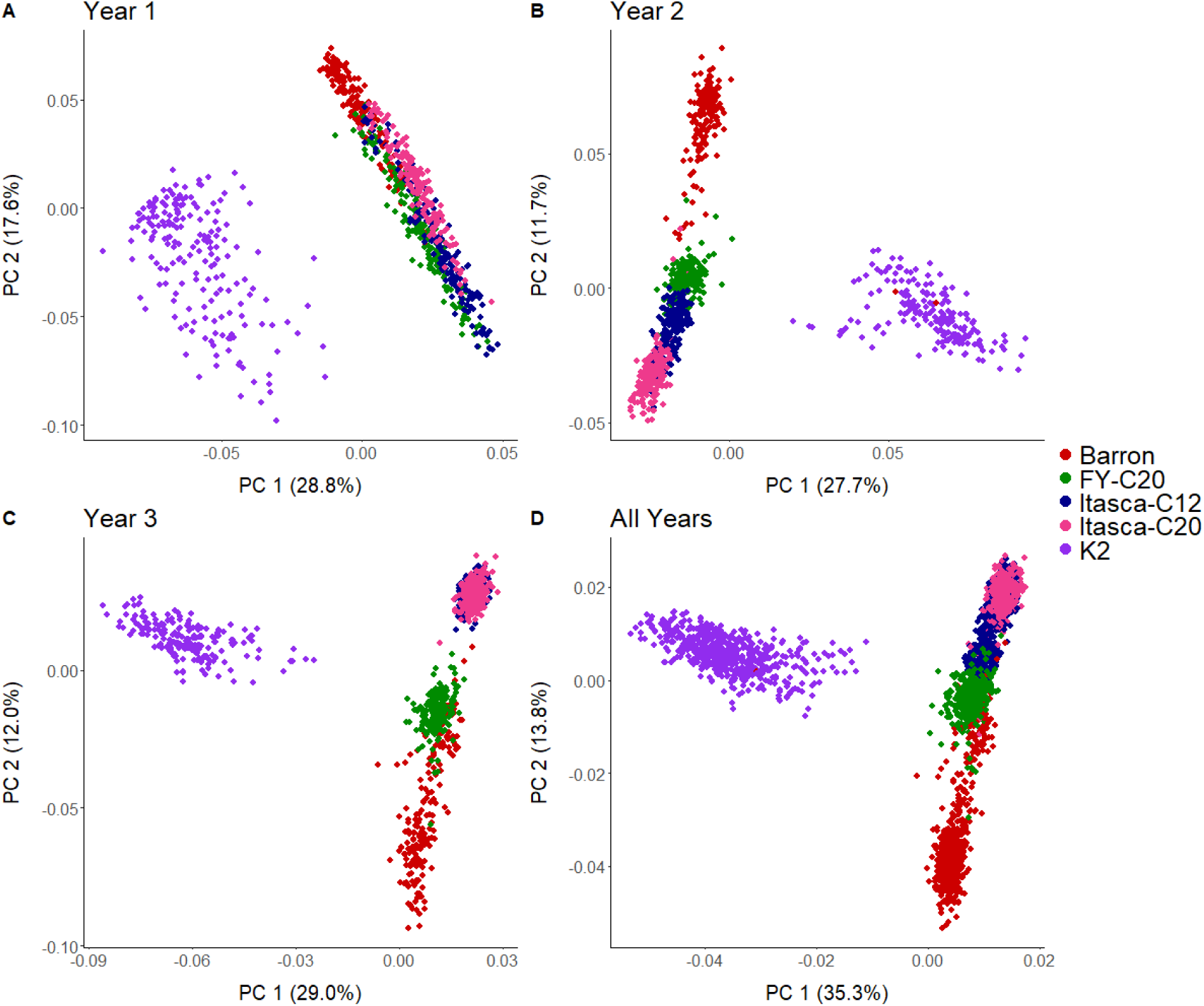
Principal component analysis (PCA) of cultivated Northern Wild Rice (cNWR; *Zizania palustris*) populations based on 73,363 single nucleotide polymorphisms (SNPs) generated via genotyping-by-sequencing (GBS). Principal components 1 and 2 (PC1 and PC2) are shown for (A) Year 1, (B) Year 2, (C) Year 3, and (D) the combined multi-year dataset. Percent variance explained by each principal component is indicated on the axes. Each point represents an individual plant, colored by population (Barron, FY-C20, Itasca-C12, Itasca-C20, and K2).

**Table 3.**
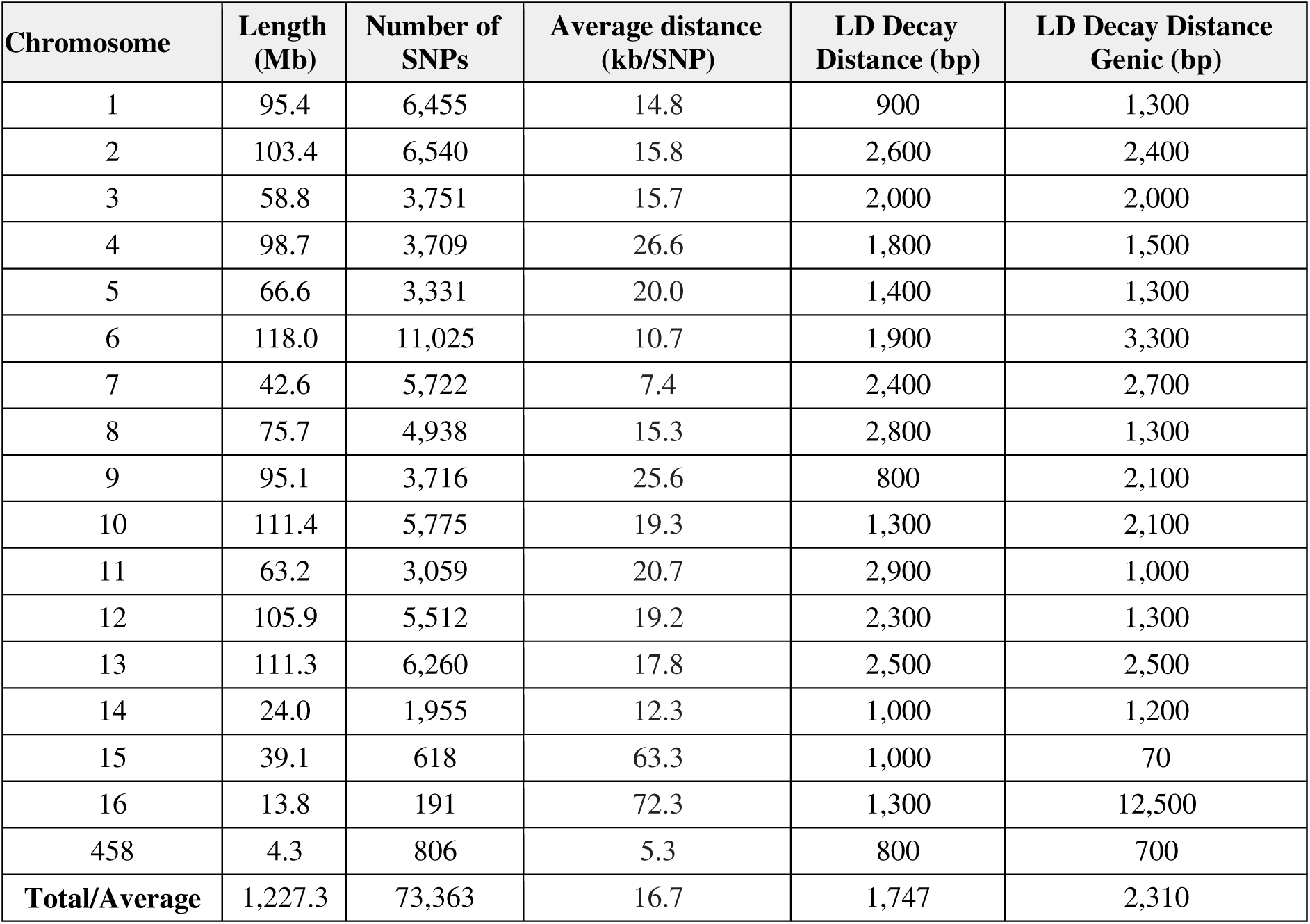
Chromosome-level summary of genome size, SNP density, and linkage disequilibrium (LD) decay in cultivated Northern Wild Rice (cNWR; *Zizania palustris*); chromosome length (Mb), number of filtered SNPs, average marker spacing (kb per SNP), and LD decay distances are shown for each chromosome. LD decay distance represents the physical distance (bp) at which pairwise LD declines to r² = 0.1, calculated using all SNPs and SNPs within annotated gene models (genic). Total/Average values represent genome-wide totals for chromosome length and SNP number and mean values for marker spacing and LD decay across chromosomes.

The pronounced separation of K2 along PC1 suggests substantial allele frequency differentiation relative to the other cultivated populations, consistent with its distinct isolation history within the breeding program (McGilp et al., 2023). In contrast, the closer clustering of Barron, the Itascas, and FY-C20 likely reflect shared ancestry and recurrent selection within partially overlapping breeding pools. A scree plot (Supplemental Figure S2) indicated that the first four PCs captured the majority of detectable structure and were therefore included in the MLM as covariates (Q matrix) to account for this stratification.

### Genome-Wide Associations

#### Overview of Association Signals

Single-plant GWAS (sp-GWAS) was conducted using a mixed linear model (MLM) with population structure and kinship included to control for relatedness and stratification. Manhattan and Q-Q plots for each trait are presented in Figure 3. No significant associations were detected for flag leaf length (FLL; Figure 3e), and subsequent analyses therefore focused on plant height (PH), primary stem width (PSW), basal stem width (BSW), and flag leaf width (FLW). Across the four traits, 124 significant SNPs (FDR < 0.01) were identified and consolidated into 98 independent loci using chromosome-specific LD decay thresholds (Table 3). Each locus consisted of one or more significant SNPs within the threshold and was treated as a distinct genomic interval associated with phenotypic variation. Of the 98 loci, 52 were trait-specific and 46 were associated with two or more traits. Fifty-six loci contained at least one significant SNP within an annotated gene model, providing biologically plausible candidates. However, given marker density and LD structure, these signals could be tagging broader genomic intervals rather than causal variants. For loci without intragenic SNPs, nearby annotated genes within the LD-defined intervals were considered as putative candidates, including potential regulatory targets. Loci were non-randomly distributed across the genome, with clustering observed on chromosomes 6, 7, 8, 10, 12, and 13 (Figure 4). Minor allele frequencies (MAF) ranged from 0.065 to 0.499 (mean = 0.284), indicating that most significant alleles segregate at moderate frequencies and may be suitable for deployment in breeding populations (Supplemental Table S6).

**Figure 3.**
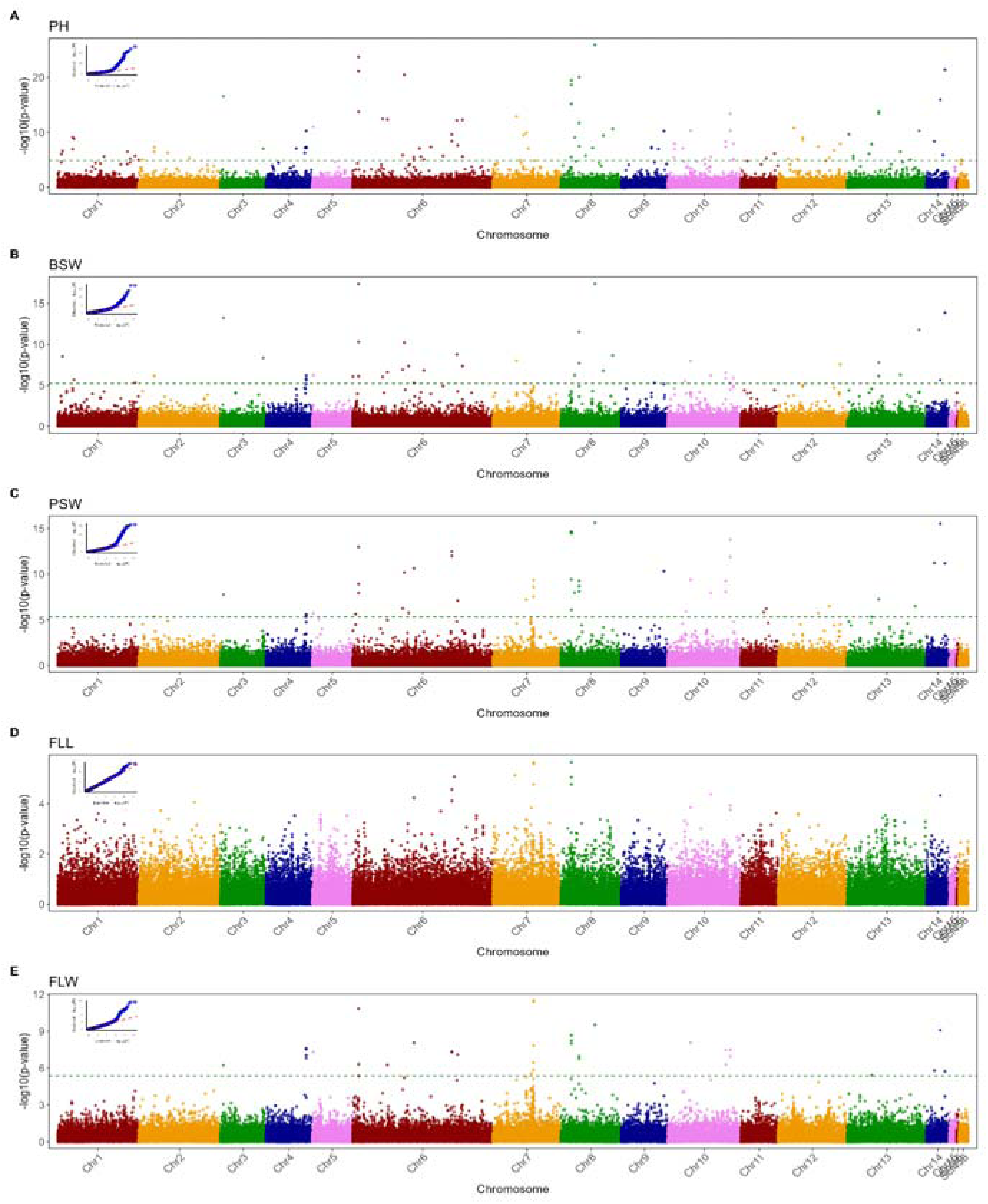
Genome-wide association analysis of vegetative architecture traits in cultivated Northern Wild Rice (*Zizania palustris* L.) represented by Manhattan plots for (A) plant height (PH), (B) basal stem width (BSW), (C) primary stem width (PSW), (D) flag leaf length (FLL), and (E) flag leaf width (FLW). Points represent 73,363 filtered SNP markers distributed across 15 chromosomes and two additional scaffolds. The x-axis shows physical position by chromosome, and the y-axis shows −logLJLJ(p-value) from the mixed linear model including population structure (first four principal components) and kinship. The horizontal dashed line indicates the false discovery rate (FDR) significance threshold of < 0.01. Quantile-quantile (Q-Q) plots inset in the upper left of each panel compare observed versus expected −logLJLJ(p-values) to assess model fit and genomic inflation.

**Figure 4.**
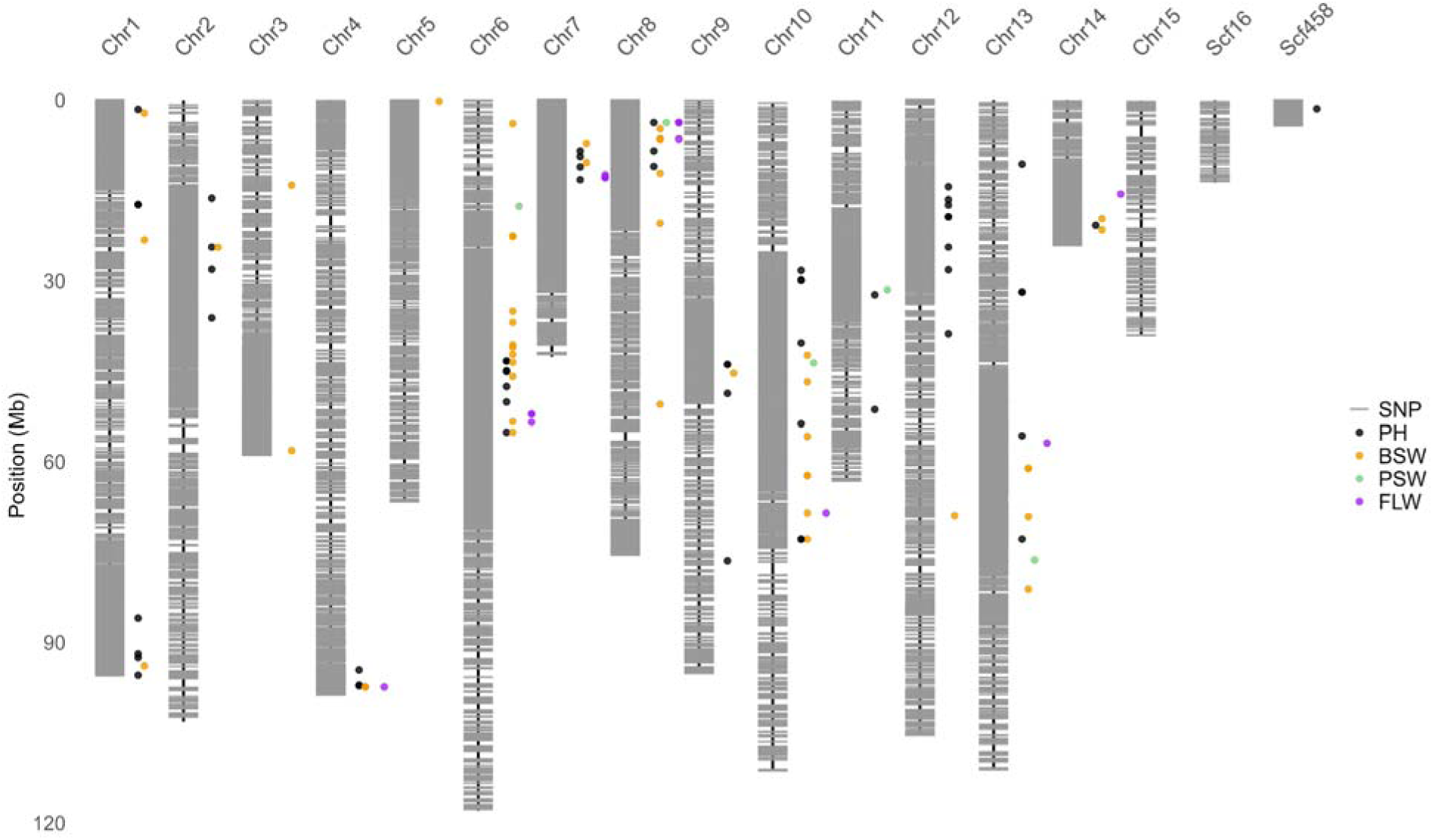
Chromosomal distribution of genome-wide SNPs and significant GWAS loci in cultivated Northern Wild Rice (*Zizania palustris* L.). Physical distribution of 73,363 filtered SNP markers and significant association signals across 15 chromosomes and two additional scaffolds of the *Z. palustris* reference genome (Haas et al., 2021). Grey vertical bars represent chromosome length (Mb), with horizontal tick marks indicating SNP density along each chromosome. Colored circles denote genomic positions of SNPs significantly associated (false discovery rate < 0.01) with vegetative architecture traits: plant height (PH, red), basal stem width (BSW, blue), primary stem width (PSW, green), and flag leaf width (FLW, orange).

#### Individual Trait Associations

Plant height (PH) exhibited the strongest genetic signal, with 85 of 98 loci associated with PH, and 43 loci unique to the trait (Figure 3a; Figure 4; Supplemental Table S6). SNP-level effect sizes ranged from −12.3 to 14.6 cm (Supplemental Table S6), consistent with polygenic control. Several loci contained biologically relevant candidates that lead to reductions in PH. For example, a locus within *ZPchr0007g5772*, encoding a putative *Tassel ear1* (*TE1*) homolog, wa associated with reduced PH (−7.33 cm). In *Zea mays*, *TE1* regulates the transition from vegetative to reproductive development, and is a key determinant of plant stature (Veit et al., 1998). Protein BLAST revealed that the predicted protein shared 65.8% amino acid identity with *ZmTE1* (NP_001104903.1, E = 0.0). A second locus associated with reduced height (−7.64 cm) was found within *ZPchr0012g20472*, a SHAQKYF-class MYB-like transcription factor, members of which function as transcriptional regulators involved in plant developmental processes and responses to environmental stress in multiple plant species(Dubos et al., 2010; Jin et al., 2017). Another signal associated with PH reduction (−5.46 cm) was within *ZPchr0013g35032*, encoding a cysteine-rich receptor-like kinase 10 (CRK10). CRK family members function in redox and immune signaling, and may exhibit pleiotropic developmental effects (Chern et al., 2016; Quezada et al., 2019). Two SNPs mapped to *ZPchr0006g44705*, encoding BRICK1 (BRK1) (−4.08 and −4.36 cm), a regulator of cytoskeletal dynamics during plant development in maize (Frank et al., 2003; Frank & Smith, 2002). Defense-associated loci were also identified, including *ZPchr0007g3720* (PYRICULARIA ORYZAE RESISTANCE 21-like; −0.11 cm) and *ZPchr0458g22510* (NBS-LRR-like resistance protein; −5.3 cm), suggesting potential growth-defense tradeoffs contributing to height variation.

Basal (BSW) and primary stem width (PSW) were associated with 39 and 35 loci, respectively, with signal enrichment on chromosomes 6, 8, and 10 (Figure 3b-c; Figure 4). Effect sizes ranged from −0.9 to 1.2 mm for BSW and −0.2 to 0.32 mm for PSW. BSW-specific loci (n = 3) were located within *ZPchr0006g44849*, encoding an aminotransferase (+0.58 mm); *ZPchr0009g576*, annotated as a β-1,3-glucosidase (−0.49 mm); and *ZPchr0010g11286*, predicted to encode a chloroplast membrane anchor (−0.56 mm). This predicted chloroplast anchor shared 89% amino acid identity (E = 0.0) with an *O. sativa* translocase of chloroplast 101 (XP_01563922.1). Four PSW-specific signals were identified within *ZPchr0010g10484*, encoding a phosphatidylinositol 3- and 4-kinase (+0.22 mm); *ZPchr0011g27442*, a pectinesterase (+0.25 mm); *ZPchr0013g33897*, a calmodulin-binding transcription activator (+0.37 mm); and one intergenic locus (+0.23 mm).

FLW exhibited the fewest associations, with 22 loci identified (Figure 3de; Figure 3) and effect sizes ranging from −0.2 to 0.2 cm. Two loci were unique to FLW. One occurred within *ZPchr0007g3720* (uncharacterized protein; effect size = −0.11 cm), while another consisted of two SNPs within *ZPchr0007g4798*, encoding a FLOWERING-PROMOTING FACTOR 1-LIKE (FPF1) protein (−0.13 cm each). Protein BLAST analysis of *ZPchr0007g4798* against *Z. mays* FPF1 protein 5 (XP_020404756.1) showed 77.0% amino acid identity (E = 2 × 10O46). In *Arabidopsis thaliana*, FPF1 promotes flowering downstream of gibberellin signaling (Kania et al., 1997), suggesting potential links between vegetative morphology and developmental phase transitions.

#### Multi-Trait Associations

Nearly half of all detected loci (46 of 98) were associated with two or more traits (Figure 3; Figure 4; Table 4), consistent with the strong positive phenotypic correlations observed among the traits (r = 0.50-0.75) (Supplemental Table S5). Eleven loci were associated with all four traits. The most prominent signal, locus 4.4, was composed of four SNPs within *ZPchr0004g39193* (*SORBI_3006G250400-like*) and was associated with parallel reductions in PH, PSW, BSW, and FLW. Notably, *ZPchr0004g39193* is directly adjacent to a *bHLH/IBH1*-like transcription factor (*ZPchr0004g39205*). In plants, *bHLH/IBH1*-like transcription factors repress cell elongation by antagonizing brassinosteroid-induced PRE transcription factors, thereby limiting internode and lamina joint elongation ( Zhang et al., 2009; Zhiponova et al., 2014). Two additional four-trait hubs, loci 6.3 and 8.3, each consisted of three SNPs associated with coordinated reductions across traits. Locus 6.3 was near *ZPchr0006g45499* (UreD urease accessory protein) and locus 8.3 was identified within *ZPchr0008g13950* (Hsp20/α-crystallin family). Notably, this locus lies immediately adjacent to *ZPchr0008g12829*, which encodes STRUCTURE-SPECIFIC RECOGNITION PROTEIN 1 (SSRP1), a subunit of the FACT (Facilitates Chromatin Transcription) chromatin remodeling complex required for normal plant development in *Arabidopsis* (Duroux et al., 2004; Lolas et al., 2010).

**Table 4.** Multi-trait loci associated with vegetative architecture in cultivated Northern Wild Rice (cNWR; *Zizania palustris*); significant SNPs (false discovery rate < 0.01) were identified using single-plant genome-wide association analysis and grouped into loci based on chromosome-specific linkage disequilibrium (LD) decay thresholds. SNP_ID denotes marker name, and chromosome (Chr.) and position refer to physical coordinates on the *Z. palustris* reference genome v1.0. Major and minor alleles indicate allelic states in the population, and MAF is the minor allele frequency. Traits include plant height (PH), flag leaf width (FLW), primary stem width (PSW), and basal stem width (BSW), with “All” indicating association with all four traits. Effect sizes represent the estimated additive effect of the minor allele (cm for PH; mm for FLW, PSW, and BSW). “Within gene” indicates whether the SNP falls within an annotated gene model; gene candidate and annotation correspond to the nearest or overlapping gene within the LD-defined interval.

| Locus | SNP_ID | Chr. | Position (bp) | Major Allele | Minor Allele | MAF | Traits | PH effect (cm) | FLW effect (mm) | PSW effect (mm) | BSW effect (mm) | Within gene | Gene Candidate | Gene Candidate Annotation |
| --- | --- | --- | --- | --- | --- | --- | --- | --- | --- | --- | --- | --- | --- | --- |
| 3.1 | SZPCHR0003_14078424 | 3 | 14,078,424 | A | C | 0.36 | All | -9.15 | -0.11 | -0.27 | -0.82 | no | - | - |
| 4.4 | SZPCHR0004_97395087 | 4 | 97,395,087 | G | A | 0.26 | All | -5.88 | -0.13 | -0.23 | -0.52 | yes | ZPchr0004g39193 | hypothetical protein SORBI_3006G250400 |
|  | SZPCHR0004_97395113 |  | 97,395,113 | C | T | 0.28 | All | -7.05 | -0.12 | -0.22 | -0.54 | yes | ZPchr0004g39193 | hypothetical protein SORBI_3006G250400 |
|  | SZPCHR0004_97395086 |  | 97,395,086 | A | C | 0.26 | PH, FLW, BSW | -5.91 | -0.13 | - | -0.53 | yes | ZPchr0004g39193 | hypothetical protein SORBI_3006G250400 |
|  | SZPCHR0004_97395047 |  | 97,395,047 | A | G | 0.27 | PH, FLW, PSW | -5.85 | -0.13 | -0.23 | - | yes | ZPchr0004g39193 | hypothetical protein SORBI_3006G250400 |
| 5.1 | SZPCHR0005_257908 | 5 | 257,908 | T | C | 0.33 | All | -7.68 | -0.13 | -0.24 | -0.57 | no | - | - |
| 6.3 | SZPCHR0006_22592445 | 6 | 22,592,445 | G | C | 0.47 | All | -9.75 | -0.14 | -0.31 | -0.83 | no | ZPchr0006g45499 | urease accessory protein D-like |
|  | SZPCHR0006_22592447 |  | 22,592,447 | A | C | 0.34 | All | -12.31 | -0.14 | -0.34 | -0.84 | no | ZPchr0006g45499 | urease accessory protein D-like |
|  | SZPCHR0006_22592448 |  | 22,592,448 | A | C | 0.25 | All | -9.01 | -0.12 | -0.30 | -0.59 | no | ZPchr0006g45499 | urease accessory protein D-like |
| 8.3 | SZPCHR0008_6458846 | 8 | 6,458,846 | T | A | 0.41 | All | -9.19 | -0.15 | -0.35 | -0.75 | yes | ZPchr0008g13950 | Hsp20/alpha crystallin family |
|  | SZPCHR0008_6458874 |  | 6,458,874 | C | A | 0.32 | PH, PSW, BSW | -11.77 | - | -0.32 | -0.89 | yes | ZPchr0008g13950 | Hsp20/alpha crystallin family |
|  | SZPCHR0008_6458853 |  | 6,458,853 | T | A | 0.24 | PH, FLW, PSW | -6.08 | -0.13 | -0.31 | - | yes | ZPchr0008g13950 | Hsp20/alpha crystallin family |
| 8.6 | SZPCHR0008_12118505 | 8 | 12,118,505 | T | G | 0.33 | All | 14.62 | 0.18 | 0.49 | 1.19 | no | ZPchr0008g11432 | OSJNBA0035M09.9 PROTEIN |
| 10.10 | SZPCHR0010_68556229 | 10 | 68,556,229 | C | T | 0.36 | All | 12.22 | 0.19 | 0.54 | 0.74 | yes | ZPchr0010g10223 | uncharacterized protein LOC4348182 |
|  | SZPCHR0010_68556238 |  | 68,556,238 | G | A | 0.37 | PH, FLW, PSW | 10.45 | 0.18 | 0.50 | - | yes | ZPchr0010g10223 | uncharacterized protein LOC4348182 |
| 10.6 | SZPCHR0010_46805763 | 10 | 46,805,763 | A | G | 0.28 | All | 7.95 | 0.15 | 0.34 | 0.70 | yes | ZPchr0010g9230 | protein G1-like9/ ALOG domain |
| 10.9 | SZPCHR0010_62282220 | 10 | 62,282,220 | T | C | 0.32 | All | 4.88 | 0.10 | 0.23 | 0.46 | no | - | - |
|  | SZPCHR0010_62282240 |  | 62,282,240 | G | T | 0.31 | All | 7.17 | 0.15 | 0.34 | 0.60 | no | - | - |
| 14.2 | SZPCHR0014_19671148 | 10 | 19,671,148 | G | T | 0.32 | All | -6.97 | -0.11 | -0.29 | -0.39 | yes | ZPchr0014g46879 | shikimate O-hydroxycinnamoyltransferase |
| 14.4 | SZPCHR0014_21522874 | 10 | 21,522,874 | C | A | 0.26 | All | 10.39 | 0.11 | 0.33 | 0.83 | yes | ZPchr0014g47155 | - |

The remaining four-trait loci were composed of one or two significant SNPs and frequently showed parallel directional effects across traits (Figure 3; Table 4). These included loci within *ZPchr0010g10223* (LOC4348182), *ZPchr0010g9230* (ALOG domain), *ZPchr0014g46879* (shikimate O-hydroxycinnamoyltransferase), as well as *ZPchr0014g47155*, which is uncharacterized but directly adjacent to *ZPchr0014g47530*. This gene encodes an elongation factor *G-1*, which is involved in chloroplast development (Ruppel & Hangarter, 2007; Tiller & Bock, 2014; Y. Zhang et al., 2023). Additional loci were intergenic but located near biologically relevant candidates. For example, locus 3.1 occurred near *ZPchr0003g18218*, encoding a zinc finger protein *GIS3* (GLABROUS INFLORESCENCE STEMS 3) implicated in gibberellin-mediated development (Gan et al., 2007). This predicted protein shared 66.89% amino acid similarity (E-value = 6 x 10^-77^) with the *O. sativa GIS3* homolog (XP_015628802.1). Locus 5.1, was downstream of *ZPchr0005g14570*, encoding the RING-type E3 ubiquitin ligase RDUF2, which modulates ABA signaling and stress-responsive growth regulation in Arabidopsis (Kim et al., 2012; Stone, 2014).

#### Multi-Trait Subsets

The remaining multi-trait loci (n = 35) formed biologically coherent subsets (Figure 4; Supplemental Table S6). A structural growth subset linking PH, PSW, and BSW (n = 21) exhibited coordinated effects on plant height and stem robustness, mirroring the moderate-to-strong correlations among PH, PSW, and BSW (r = 0.55-0.60). Locus 2.3 was located within *ZPchr0002g23763*, annotated as ROOT PRIMORDIUM DEFECTIVE 1 (RPD1), a nuclear protein required for the initiation of root primordia and normal meristematic development in *Arabidopsis* (Konishi & Sugiyama, 2006). Variation in this gene was associated with coordinated reductions in PH (−6.36 cm), PSW (−0.23 mm), and BSW (−0.56 mm). The predicted protein shared 85.21% amino acid identity (E = 0.0) with a *Panicum miliaceum* homolog (RLN34983.1). Additional signals included *ZPchr0006g41703* (microtubule-associated protein) and *ZPchr0006g41107* (a neurochondrin-like protein), proteins that participate in cytoskeletal organization and cell differentiation processes influencing stem elongation and mechanical strength during vegetative development (Sedbrook & Kaloriti, 2008).

The canopy growth subset (PH-FLW-PSW, n = 11) connected vertical growth with upper-canopy morphology. The inclusion of FLW within this subset is consistent with its strong correlation with PSW (r = 0.75) and moderate correlation with PH (r = 0.50). Locus 8.1 within *ZPchr0008g13061* (*bHLH13* transcription factor) was associated with reduced PH and PSW. Locus 6.18 was located in *ZPchr0006g42764*, encoding a squamosa promoter-binding-like (*SPL*) transcription factor and was associated with increased PH and FLW (Figure 4; Supplemental Table S6). The predicted protein shared 66.7% amino acid identity with *O. sativa OsSPL14/IPA1* (LOC_Os08g39890; NP_001064623.1, E = 1 × 10D32), a regulator of vegetative phase transition and plant architecture (Jiao et al., 2010; Miura et al., 2010).

### Candidate Gene and Diplotype Analysis

To further investigate the genomic regions underlying significant GWAS associations, we conducted fine-scale candidate gene and diplotype analyses for six loci, with loci 4.4 and 8.1 highlighted here as representative examples of contrasting local genetic architecture. This approach integrated local sequence variation, gene expression data, haplotype structure, and phenotypic distributions to evaluate whether GWAS signals reflect broader regional genetic effects rather than single-marker associations. Using a cNWR gene expression atlas (Banting et al., 2026), tissue-specific expression profiles were examined to assess potential functional relevance of candidate genes within. For each locus, diplotypes were defined using both significant and nearby non-significant SNPs, allowing assessment of the local haplotype and its relationship to phenotypic variation. Diplotypes with a count of over 25 individuals for loci 4.4, 6.3, 6.16, 6.18, 8.1, and 8.3 ranged from 2 to 29 in number, with an average of ∼17 diplotypes per locus (Figure 5-6, Supplemental Figures S3-S6). The diplotypes were made up from between 2 and 16 haplotypes, with an average of 11 per locus. All diplotypes were associated with distinguishable phenotypic differences with the exception of loci 6.18 (Supplemental Figure S5). Together, these examples highlight the range of genetic architectures contributing to quantitative variation in cNWR.

**Figure 5.**
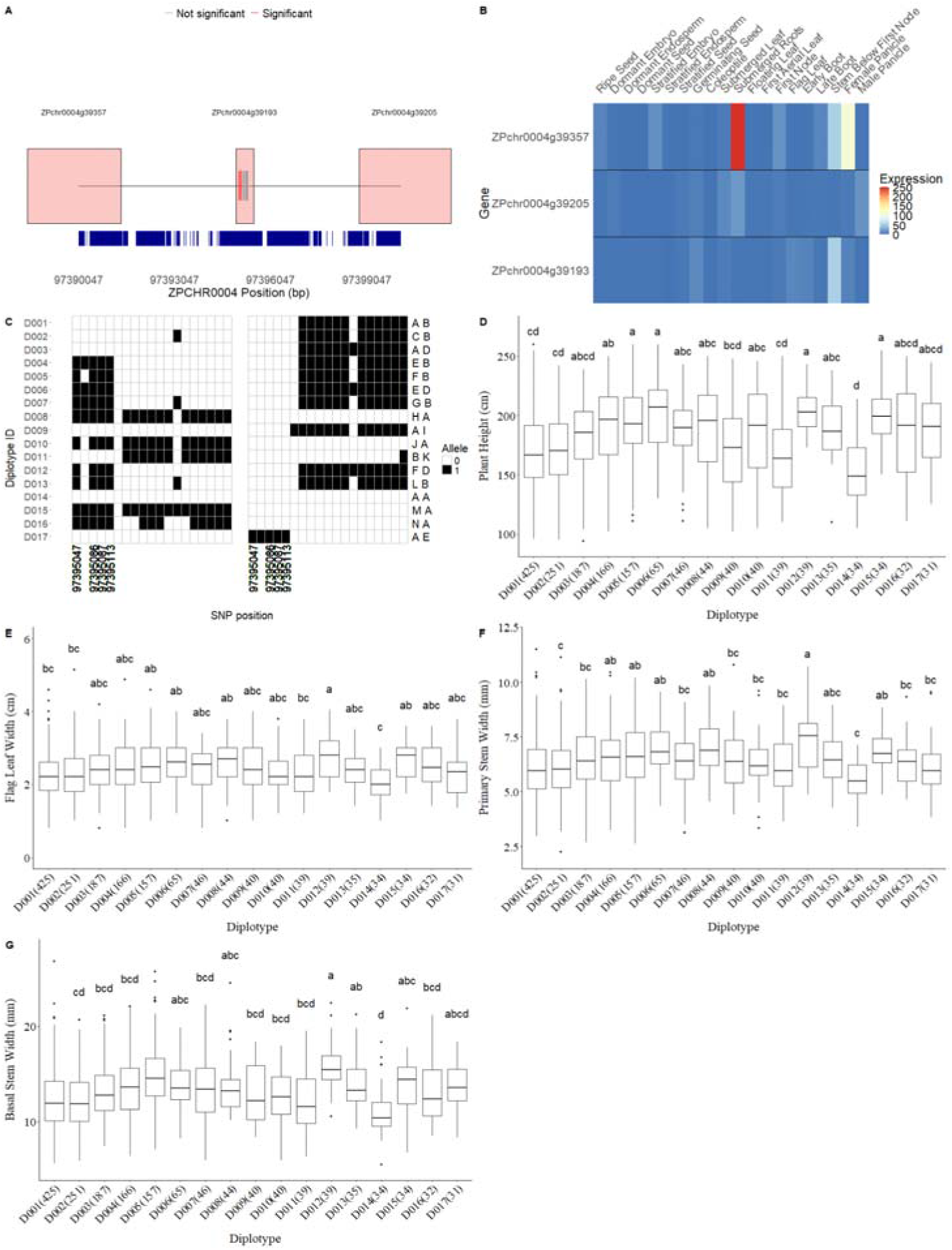
Diplotype structure and genic and phenotypic effects at locus 4.4 in cultivated Northern Wild Rice (*Zizania palustris* L.); (A) local genomic region of locus 4.4 on chromosome 4, (B) gene expression across growth stages, (C) diplotype structure inferred from phased SNPs within ±3 kb of the locus, with haplotypes labeled by capital letters, and (D–G) phenotypic distributions of diplotype classes for plant height (PH), flag leaf width (FLW), primary stem width (PSW), and basal stem width (BSW), respectively. Lowercase letters indicate significant differences among diplotypes based on Tukey’s honest significant difference (HSD) test (α = 0.05).

**Figure 6.**
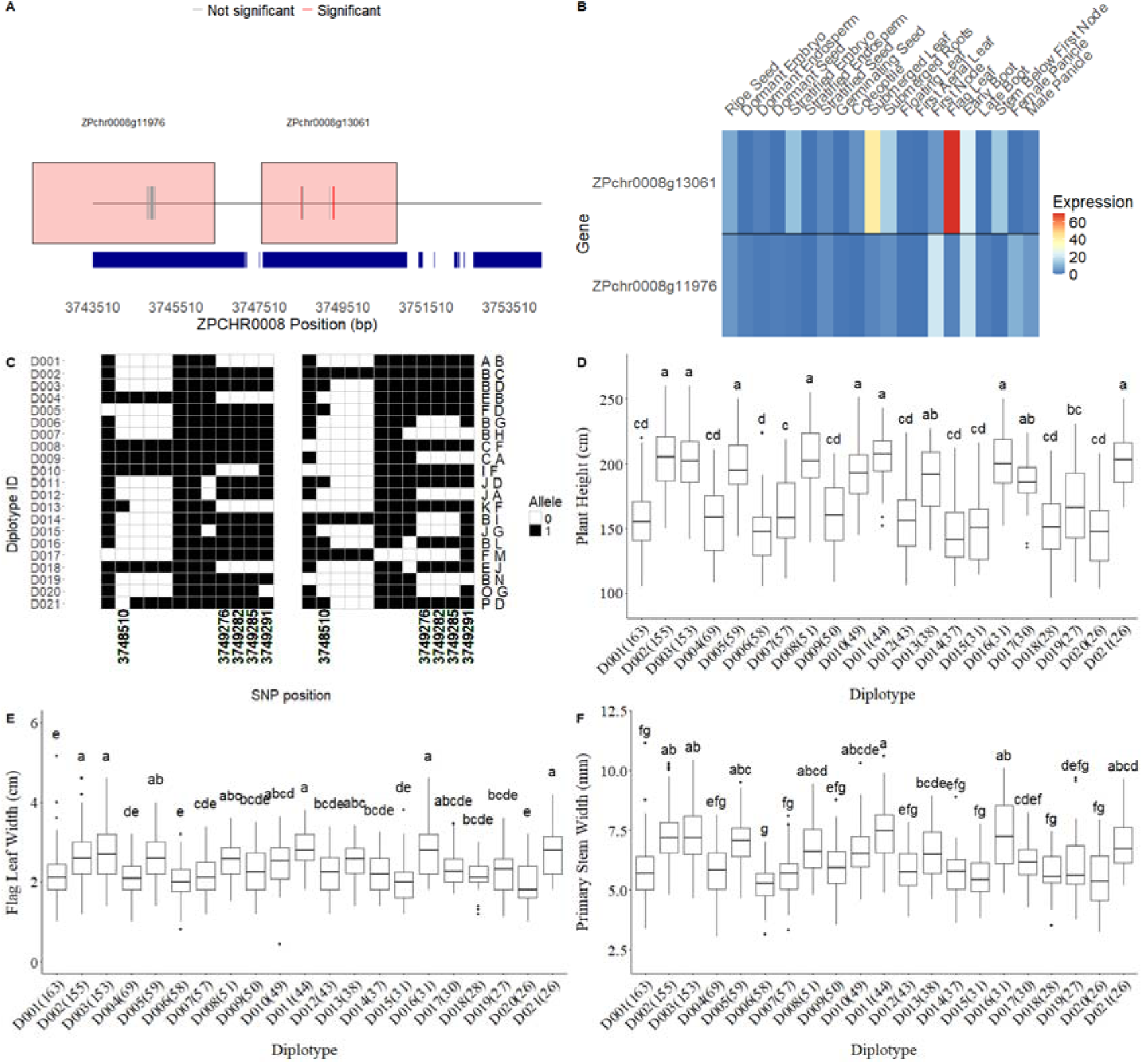
Diplotype structure genotypic and phenotypic effects at locus 8.1 in cultivated Northern Wild Rice (*Zizania palustris* L.); (A) Local genomic region of locus 8.1 on chromosome 8, (B) gene expression across growth stages C) diplotype structure inferred from phased SNPs located within ±3 kb of the locus, with haplotypes labeled by capital letters, and (D-F) phenotypic distributions of diplotype classes for (D) plant height (PH), (E) flag leaf width (FLW), and (F) primary stem width (PSW), respectively. Lowercase letters indicate significant differences among diplotypes based on Tukey’s honest significant difference (HSD) test (α = 0.05).

#### Locus 4.4

On chromosome 4, fine-mapping of locus 4.4 identified three annotated gene models within the LD-defined interval: *ZPchr0004g39357, ZPchr0004g39193,* and *ZPchr0004g39205* (Figure 5A). Four significant and fifteen non-significant SNPs were concentrated within *ZPchr0004g39193*, with no other SNPs found within the interval or other gene models. The highest expression was found for *ZPchr0004g39357*, particularly during the submerged root (∼250 Transcripts Per Million (TPM)), female panicle (∼109 TPM) and the stem below the first panicle (∼56 TPM), growth stages with all other stages having expression below 17 TPM (Figure 5B) (Banting et al., 2026). Gene expression for *ZPchr0004g39193* was highest in stem growth before the first node (∼55 TPM) but was low (<12 TPM) across all other growth stages. Expression for *ZPchr0004g39205* was highest in submerged root tissue (∼21 TPM) and male panicle tissue (∼17 TPM) but low (<12 TPM) at all other growth stages.

A total of fourteen haplotypes (A-N) were identified across the locus, forming 17 diplotype classes (Figure 5C). ANOVA and Tukey’s HSD were used to assess phenotypic differences between diplotypes for each trait. For plant height, diplotypes displayed moderate but consistent separation among HSD groupings (Figure 5D). Several diplotypes containing haplotypes such as D, J, and K were frequently observed among the taller phenotypic classes (“a” or “ab”), indicating association with increased PH across multiple haplotype pairings. In contrast, diplotypes containing haplotypes such as A or M were more frequently classified within shorter height groupings. Other haplotypes occurred in both taller and intermediate classes depending on pairing, suggesting context-dependent effects. Patterns for FLW were broadly consistent with those observed for PH (Figure 5E). Diplotypes containing haplotypes J and K were commonly observed among wider-leaf groupings, whereas diplotypes containing A or M tended to occur among narrower classes. However, substantial overlap among intermediate HSD groupings indicates that the locus contributes modestly to FLW, relative to overall phenotypic variance. PSW and BSW also showed patterns similar to those observed for PH and FLW (Figure 5F-G). Diplotypes containing haplotypes J and K were frequently associated with thicker stems, while diplotypes containing A or M were more commonly associated with thinner stems. Intermediate haplotypes appeared across multiple HSD groupings depending on pairing, again indicating context-dependent phenotypic effects. Overall, locus 4.4 exhibited coordinated effects across plant height, leaf width, and stem robustness, suggesting that allelic variation within this genomic interval contributes to multiple aspects of plant architecture.

#### Locus 8.1

On chr. 8, fine-mapping of locus 8.1 identified two annotated gene models, *ZPchr0008g11575* (putative importin-alpha re-exporter) and *ZPchr0008g13061*, within the LD-defined interval (Figure 6A). The locus encompassed 12 SNPs, five of which were significant and located within *Zpchr0008g13061*. Expression of *ZPchr0008g11575* was highest in first node tissue (∼21 TPM) and at the early boot stage (∼20 TPM) but remained below 10 TPM at all other growth stages. Gene expression for *Zpchr0008g13061* was highest during the flag leaf (∼70 TPM), submerged leaf (∼42 TPM) and early boot (∼23 TPM) growth stages, while all other stages remained below 16 TPM (Figure 6B).

Sixteen haplotypes (A-P) were identified, forming 21 diplotype classes represented by ≥25 individuals (Figure 6B). For plant height, diplotypes largely separated into two statistically distinct HSD groupings corresponding to taller (“a”) and shorter (“cd”) phenotypic classes (Figure 6C). Several haplotypes, including D and K, were repeatedly present in taller diplotypes across multiple pairings, indicating consistent association with increased mean height across diverse genetic backgrounds. In contrast, diplotypes containing haplotypes L, A, and C were consistently classified among shorter groups. Other haplotypes, including B, E, F, and G, were observed in both taller and shorter diplotype classes depending on pairing, suggesting context-dependent phenotypic effects. Haplotypes such as N and M were less frequent, limiting robust inference regarding their independent contributions.

Flag leaf width exhibited broader phenotypic dispersion and less discrete clustering than PH (Figure 6C). Nonetheless, diplotypes containing D and K haplotypes were disproportionately represented in wider-leaf HSD groupings (“a”), while diplotypes containing A or L were more frequently associated with narrower classes (“e”). Overlap among intermediate groupings indicates that locus 8.1 contributes modestly to FLW, relative to total phenotypic variance. Primary stem width (PSW) displayed patterns similar to PH and FLW (Figure 6D). Diplotypes containing GK were among the thickest-stem classes, whereas diplotypes containing A, C, I, and L were more frequently associated with thinner stems. Together, these results indicate that locus 8.1 contributes to coordinated variation in height, stem robustness, and leaf width, with effects dependent on specific haplotype combinations.

## DISCUSSION

### sp-GWAS in a Non-Replicable, Outcrossing Crop

This study represents the first genome-wide association analysis conducted in cultivated Northern Wild Rice, an obligately outcrossing crop with self-incompatibility and atypical seed storage behavior that precludes the replicated, inbred resources used in conventional GWAS. By leveraging large per-year sample sizes across three field seasons, we identified numerous loci associated with vegetative architecture traits despite low to moderate broad-sense heritabilities. The rapid LD decay observed in this panel enabled relatively fine-scale localization of association signals, with multiple significant SNPs frequently collapsing into compact, locus-level intervals under chromosome-specific LD thresholds (Browning et al., 2021; Myles et al., 2009). While causal resolution remains limited, these results indicate that sp-GWAS can recover biologically meaningful association signals in highly heterozygous, non-replicable crop systems when sample sizes are large and mixed models appropriately account for relatedness and structure. In this respect, our findings extend prior demonstrations in maize and luffa (*Luffa* spp.) to an aquatic, self-incompatible cereal system (Gyawali et al., 2019; Li et al., 2024).

### Environmental Effects and Recoverable Genetic Signal

Vegetative morphology in cNWR was strongly influenced by year-to-year environmental variation, with year effects explaining a substantial proportion of phenotypic variance across traits (6.8 - 54.1%). Broad-sense heritability estimates were correspondingly low to moderate (H² = 0.03 - 0.34), consistent with pronounced phenotypic plasticity in stem elongation, culm thickness, and leaf morphology. This pattern aligns with broader cereal literature, where architecture traits integrate multiple developmental processes and respond dynamically to temperature, resource availability, and water status (Sakamoto & Matsuoka, 2008; Weng et al., 2011). In aquatic paddy systems, additional hydrologic heterogeneity (e.g., water depth gradients, current, and within-paddy microclimates) likely amplifies environmental variance relative to upland cereal trials (Oelke & Porter, 2016).

Despite this environmental noise, we detected numerous significant loci across traits. This suggests that sufficient additive genetic signal persists in cNWR architecture to enable locus detection under large sample sizes and multi-year sampling. Importantly, the identification of loci under these conditions should not be interpreted as evidence of large-effect variants, but rather as evidence that polygenic signals remain detectable even when phenotypic variance is strongly environmentally modulated. These results support multi-year sampling as a practical framework for architecture trait dissection in cNWR and similar outcrossing systems.

### Polygenic and Integrated Architecture of Vegetative Traits

The strong and uniformly positive phenotypic correlations among vegetative traits provided a useful framework for interpreting the genetic architecture uncovered here. Across traits, association signals were distributed across many loci of small to moderate effect, consistent with a quantitative genetic architecture. Plant height exhibited the largest number of associated loci and the broadest range of estimated SNP effects, whereas stem width and flag leaf width showed fewer associations. This pattern is consistent with GWAS findings in other cereals, where plant height is among the most genetically tractable architecture traits due to its cumulative integration of cell elongation, internode development, and hormonal regulation (X. Huang et al., 2010; Sakamoto & Matsuoka, 2008; Wallace et al., 2014). In contrast, stem and leaf width traits often display greater environmental sensitivity and lower repeatability, reducing power to detect associations even when genetic control is present (Holland et al., 2002; Poorter et al., 2009). Variation in association strength among traits may therefore reflect differences in trait plasticity and measurement precision rather than differences in biological importance.

A central outcome of this study is the prevalence of overlapping association signals, with 46 of 98 loci associated with two or more traits and eleven loci associated with all four traits (PH, BSW, PSW, FLW). This genetic overlap closely mirrors the underlying phenotypic correlation structure observed in this panel (Supplemental Table 5). Stem and leaf width traits were strongly correlated (e.g., PSW-FLW r = 0.75; PSW-BSW r = 0.60), and plant height showed moderate correlations with both stem widths and flag leaf width (r = 0.50-0.59). These relationships indicate that the vegetative phenotype behaves as an integrated architectural complex rather than a set of independent traits, and the GWAS results recapitulate this phenotypic coupling at the locus level. This degree of shared association is consistent with partially coordinated developmental regulation of vegetative growth, in which stem elongation, culm thickness, and leaf expansion are not independent outcomes but components of an integrated growth program ( Huang et al., 2010; Zhang et al., 2021). Such associations reflect the modular nature of plant development, in which shared hormonal and transcriptional programs coordinate growth across multiple organs (Sakamoto & Matsuoka, 2008; Weng et al., 2011). The partitioning of multi-trait loci into a structural growth subset (PH-PSW-BSW) and a canopy growth subset (PH-PSW-FLW) further supports a modular interpretation of vegetative architecture variation in cNWR.

Notably, no significant loci were detected for flag leaf length in this panel. In contrast to the strong correlations observed among PH, PSW, BSW, and FLW, FLL exhibited comparatively weak associations with other traits (e.g., PH-FLL r = 0.22), suggesting partial developmental independence from the primary architecture axis. The absence of GWAS signals for FLL therefore aligns with its weaker integration into the correlated vegetative module detected for the other traits. Given the moderate heritability estimated for FLL, this result may also reflect a combination of highly polygenic control, limited marker-trait linkage at the available marker density, and/or reduced statistical power under single-plant phenotyping conditions.

### Conserved Regulatory Classes and Candidate Genes

Although GWAS associations do not demonstrate causality, the candidate gene regions identified in this study were enriched for regulatory gene classes implicated in vegetative architecture across grasses, which is consistent with the strong multi-trait signal observed in this study. Across the 98 loci detected (46 multi-trait; 11 shared by all four traits), many intervals contained transcriptional regulators, signaling components, and chromatin-associated factors.

A notable height-associated signal occurred within a putative *TE1* homolog (*ZPchr0007g5772*). In maize, *TE1* encodes a pleiotropic RNA-binding protein that regulates shoot apical and intercalary meristem activity, influencing internode elongation, leaf initiation rate, and inflorescence architecture (Veit et al., 1998; Wang et al., 2021). Expression profiling in cNWR showed strong enrichment of *ZPchr0007g5772* transcripts in dormant seed (∼1049 Transcripts Per Million (TPM)) and reproductive tissues, including male panicles (∼448 TPM) and female panicles (∼85 TPM), with additional expression detected in subterranean root tissue (∼26 TPM) and nodal tissue (∼6 TPM) (Banting et al., 2026). The presence of transcripts in dormant seed may reflect stored mRNA associated with early embryonic developmental programs, while the strong expression observed in panicle tissues is consistent with a role in meristematic activity and inflorescence development. The recovery of a *TE1*-like candidate in cNWR therefore supports conserved contributions of meristem regulation to present variation in plant height within Oryzeae and provides a mechanistically coherent hypothesis for follow-up (e.g., stage-specific expression during elongation and allele-specific expression in heterozygotes).

Signals implicating HLH/bHLH growth regulation were also consistent with conserved cereal pathways. Multi-trait hubs and canopy-linked loci occurred within or adjacent to HLH/bHLH candidates, including locus 8.1 (annotated as a bHLH transcription factor) and locus 4.4 (located adjacent to an IBH1-like factor; *ZPchr0004g39205*). In rice and Arabidopsis, the antagonistic HLH/bHLH module involving ILI1/PRE1 and IBH1 mediates brassinosteroid-linked control of cell elongation and plant development, with IBH1 acting as a growth repressor and ILI1/PRE1 counteracting this inhibition through protein interaction (Zhang et al., 2009). Expression profiles of the cNWR candidates are broadly consistent with roles in growth-associated tissues. The locus-associated gene *ZPchr0004g39193* showed strongest expression in nodal tissue and flag leaves, with additional lower expression in early boot stages and female panicles (Figure 5B) (Banting et al., 2026), suggesting activity in structural tissues associated with internode and canopy development. The adjacent IBH1-like candidate *ZPchr0004g39205* was expressed in subterranean root, submerged leaf, and male panicles (Banting et al., 2026), consistent with regulatory activity across actively growing vegetative and reproductive organs. The coordinated directional effects across plant height, stem width, and flag leaf width observed for several multi-trait loci in cNWR are therefore consistent with quantitative modulation of elongation programs operating across multiple tissues.

Other candidates to explore include a *SPL*-class transcription factor candidate (*ZPchr0006g42764*) homologous to *OsSPL14/IPA1*, a central node in the rice “ideal plant architecture” pathway regulated by miR156. In rice, allelic variation affecting *OsSPL14* regulation produces major changes in architecture and lodging-related traits (Jiao et al., 2010; Miura et al., 2010; J. Wang et al., 2017). We also identified a FLW-specific association within an FPF1-like gene, which is consistent with literature indicating that these small proteins participate in flowering-related pathways and can affect stem growth during the reproductive transition, though functional directionality can differ among grasses and model systems (Liu et al., 2023; Takagi et al., 2025).

Taken together, these candidates may indicate that quantitative architectural variation in cNWR arises from the modulation of multiple conserved regulatory nodes governing elongation, tissue differentiation, and developmental timing. Rather than being driven by a small number of large-effect, lineage-specific mutations, the observed architecture likely reflects polygenic tuning of shared transcriptional and hormonal networks. At the same time, many associated loci were intergenic or lacked obvious coding candidates within LD-defined intervals, underscoring the probable importance of regulatory variation outside annotated gene bodies. For example, *ZmTE1* expression is influenced by cis-regulatory elements located more than 1 kb from the coding sequence, illustrating how noncoding variation may modulate meristem activity and plant stature without altering protein structure (e.g., Wang et al., 2021). Future work integrating developmental-stage expression profiling, allele-specific expression analyses, and functional assays will be required to distinguish direct regulatory effects from linked associations and to clarify how these conserved gene families contribute to phenotypic diversity in cNWR.

### Diplotype-Based Interpretation of Locus Effects in an Obligate Outcrosser

The present study demonstrates that plant architectural variation in cNWR is influenced by genomic regions exhibiting distinct haplotype architectures consistent with expectations for an obligately outcrossing species. Because individuals carry two potentially different haplotypes at each locus, phenotypes reflect the combined effects of both alleles rather than simple Mendelian dominance relationships. In this context, diplotype-based analyses provide a biologically appropriate framework for interpreting GWAS signals in highly heterozygous populations and allow evaluation of how local haplotype combinations contribute to phenotypic variation.

The value of this approach was evident in the architectures of loci 8.1 and 4.4. Locus 8.1 exhibited high haplotype diversity and complex diplotype structure, with sixteen haplotypes forming twenty-one diplotype classes. Phenotypic stratification for plant height occurred across many haplotype combinations, with several haplotypes (e.g., D and K) consistently associated with taller plants across multiple genetic backgrounds, whereas others (e.g., A, C, and L) were repeatedly associated with shorter phenotypes. These patterns are consistent with a multi-allelic locus in which functional variation is represented by haplotype backgrounds rather than individual SNPs. Similarly, locus 4.4 displayed 14 haplotypes forming 17 diplotype classes showing consistent directional patterns across plant height, flag leaf width, and stem widths. Haplotypes such as J and K were repeatedly associated with taller plants and thicker stems, whereas haplotypes such as A and M were consistently associated with reduced plant stature and narrower leaves. The coordinated effects across multiple architectural traits suggest that variation within this genomic interval contributes to integrated regulation of canopy development.

Fine-scale mapping of locus 4.4 revealed a highly localized pattern of detected sequence variation. All significant and nearby non-significant SNPs occurred within the annotated gene model *ZPchr0004g39193*, while no SNPs were detected within the adjacent gene models in this interval, including the neighboring bHLH transcription factor gene (*ZPchr0004g39205*). The absence of SNPs within the neighboring bHLH gene model likely reflects the distribution of detectable polymorphism in the current dataset, not evidence that the region lacks underlying variation. Limited marker coverage in some genomic regions is expected in reduced-representation genotyping approaches, and additional resequencing would be required to fully evaluate variation across the interval.

These diplotype results also align with expectations given the rapid LD decay observed in this panel. When LD blocks are short, functional variation may be distributed across a small region (regulatory and coding variants), and haplotype-based markers can better represent the causal segment than any single SNP. Recent plant-focused frameworks explicitly highlight haplotype blocks as useful units for fine-mapping and trait dissection in complex trait architectures (Wu et al., 2022) and for defining deployable superior haplotypes in breeding programs (Bhat et al., 2021).

### Breeding Implications and Future Perspectives

The genetic architecture revealed here has several direct implications for improvement of cNWR. First, vegetative architecture traits in this panel exhibited low to moderate broad-sense heritability and substantial year effects, with environmental variance explaining up to 54% of phenotypic variation. Under such conditions, response to phenotypic selection is expected to be slow and inconsistent, particularly when selection decisions are made on single-plant observations in heterogeneous paddy environments. Similar limitations have motivated the adoption of genomic prediction in other crops for traits characterized by moderate heritability and strong genotype-by-environment interaction (Crossa et al., 2017; Heffner et al., 2009).

Second, the prevalence of multi-trait loci indicates that vegetative architecture in cNWR is genetically integrated. Direct selection on plant height alone is therefore likely to induce predictable correlated responses in stem robustness and canopy morphology, given the moderate correlations between PH and stem width traits (r ≈ 0.55-0.59) and the strong correlation between PSW and FLW (r = 0.75). However, the direction of desirable change may not always align across traits; for example, breeding for reduced plant height while maintaining or increasing stem thickness. These trade-offs highlight the importance of accounting for trait correlations in selection strategies. Multi-trait genomic selection models can explicitly leverage this genetic covariance to improve prediction accuracy relative to single-trait models when traits are genetically correlated (Calus & Veerkamp, 2011; Jia & Jannink, 2012).

These results support evaluating prediction-based selection as a complement or alternative to direct phenotypic screening. When replication is biologically constrained, genomic selection can recover additive genetic signals by aggregating small effects genome-wide and by borrowing information from across related individuals and environments (Crossa et al., 2017; Heffner et al., 2009). The distributed nature of associations detected here, particularly for plant height, further supports a polygenic architecture better suited to prediction-based breeding than to marker-assisted selection targeting a small number of QTL. The diplotype analyses of locus 8.1 supports this conclusion as the locus displayed extensive haplotype diversity and non-additive effects across diplotype combinations, indicating that functional variation is likely distributed across multiple linked polymorphisms. Such multi-allelic locus structure complicates single-SNP deployment but aligns well with genomic prediction frameworks that incorporate haplotype or genome-wide marker effects simultaneously. Recent work in multiple crop systems demonstrates that haplotype-based predictors can enhance genomic prediction accuracy when LD blocks are short and functional variation is multi-allelic (Bhat et al., 2021; Wu et al., 2022).

Finally, the identification of conserved regulatory classes, including *TE1*-like meristem regulators, BR-responsive *HLH/bHLH* factors, *SPL*-class transcription factors, and chromatin-associated components, provides biologically interpretable anchors for integrating functional genomics with breeding. Although these loci do not fall in known domestication genes, they suggest developmental modules that can be monitored through targeted sequencing, expression analysis, or haplotype tracking as breeding populations advance. Over time, coupling genomic prediction with mechanistic insight into conserved regulatory pathways may enable breeders to shift architecture toward ideotypes optimized for aquatic cultivation, balancing height, stem strength, canopy efficiency, and harvestability.

In summary, the polygenic, coordinated, and environmentally responsive architecture uncovered in this study strongly favors genomic prediction and multi-trait selection strategies over single-marker or single-trait selection approaches. cNWR thus represents both a practical case study in applying genomic selection to an obligately outcrossing cereal and a broader model for genomic improvement in emerging, partially domesticated crops.

## CONCLUSIONS

Single-plant GWAS identified statistically robust associations for vegetative architecture traits in cultivated Northern Wild Rice despite the biological constraints of an obligately outcrossing, self-incompatible crop and the strong environmental influence observed across years. Across 2,173 plants evaluated over three seasons, 98 loci were associated with plant height, stem width, and flag leaf width, indicating that useful genetic signals can be recovered in cNWR when analytical approaches are matched to species biology.

The results support a model in which vegetative architecture in cNWR is polygenic and developmentally integrated. Nearly half of detected loci were associated with multiple traits, consistent with the observed phenotypic correlations among plant height, stem width, and flag leaf width. Candidate intervals frequently contained genes belonging to conserved regulatory classes implicated in grass development, including TE1-like meristem regulators, HLH/bHLH transcription factors, SPL-class developmental regulators, and chromatin-associated components. At the same time, diplotype analyses showed that several GWAS signals are more appropriately interpreted as local haplotype effects than as isolated SNP associations, which is consistent with rapid LD decay and the highly heterozygous nature of this crop.

Together, these findings establish a foundation for genome-enabled improvement of cNWR and support future evaluation of multi-trait genomic prediction approaches for plant architecture in aquatic production systems. More broadly, this study highlights the usefulness of sp-GWAS for genetic analysis in emerging crop species where replicated inbred mapping populations are not feasible.

## SUPPLEMENTAL MATERIALS

Supplemental tables include information on trait definitions, number of plants tested, weather data, phenotypic means by year, trait correlations and a full list of significant SNPs found in the study. Supplemental sfigures include a PCA with more PCs, a scree plot showing variance by PC and gene and haplotype models for additional loci of interest.

## DATA AVAILABILITY

Raw sequence reads are available on the sequence read archive (SRA) under BioProject ID PRJNA1428252 (<u>Reviewer Link</u>). A sample key, phenotype file and filtered VCF are available on DRUM (Submitted; currently being approved). Code is available on GitHub (Release upon acceptance of article).

## AUTHOR CONTRIBUTIONS

**Lillian McGilp** Formal analysis, Data Curation,Visualization, Writing-Original Draft, Writing-Review and Editing; **Reneth Millas** Investigation, Methodology, Data Curation, Writing-Original Draft; **Matthew Haas** Data Curation, Methodology; **Alan Mickelson** Conceptualization, Investigation, Project Administration; **Laura Shannon** Supervision, Writing-Review and Editing; **Jennifer Kimball** Conceptualization, Funding Acquisition, Methodology, Project Administration, Writing-Original Draft, Writing-Review and Editing

## ABBREVIATIONS

BSW: basal stem width
BWA-MEM: burrows-wheeler aligner maximal exact match aligner
CDD: conserved domain database
cNWR: cultivated northern wild rice
FDR: false discovery rate
FLL: flag leaf length
FLW: flag leaf width
GBS: genotype-by-sequencing
GWAS: genome-wide association study
HSD: tukey’s honestly significant difference
LD: linkage disequilibrium
MLM: mixed linear model
NWR: northern wild rice
PC: principal component
PCA: principal component analysis
PH: plant height
PSW: primary stem width
QTL: quantitative trait locus
SNP: single nucleotide polymorphism
sp-GWAS: single-plant genome-wide association study
TPM: transcripts per million
UMGC: university of minnesota genomics center
UMN: university of minnesota
VCF: variant call format

## SUPPLEMENTAL TABLES AND FIGURES

**Supplemental Table S1.** Trait abbreviations, units and descriptions of measurement method for sp-GWAS in northern wild rice (*Zizania palustris* L.)

| **Trait** | **Trait abbreviation** | **Unit** | **Measurement** |
| --- | --- | --- | --- |
| Plant Height | PH | cm | The length from the soil line to the tip of the female panicle |
| Basal Stem Width | BSW | mm | The width of the base of the primary stem |
| Primary Stem Width | PSW | mm | The width of the primary stem at the flag leaf |
| Flag Leaf Length | FLL | cm | The length from the base of the flag leaf to its tip |
| Flag Leaf Width | FLW | cm | The width of the flag leaf at its widest point |

**Supplemental Table S2.** Number of individuals from each cultivated Northern Wild Rice (cNWR; *Zizania palustris*) population, collected across three years, used in genome-wide association analyses.

| Trait | Year | Barron | FY-C20 | Itasca-C12 | Itasca-C20 | K2 | Total |
| --- | --- | --- | --- | --- | --- | --- | --- |
| Plant Height (PH) | 1 | 145 | 148 | 119 | 145 | 180 | 737 |
|  | 2 | 165 | 180 | 175 | 199 | 159 | 878 |
|  | 3 | 196 | 190 | 197 | 200 | 198 | 981 |
| Total |  |  |  |  |  |  | 2596 |
| Basal Stem Width (BSW) | 1 | 146 | 156 | 155 | 155 | 177 | 789 |
|  | 2 | 166 | 180 | 176 | 198 | 186 | 906 |
|  | 3 | 200 | 189 | 199 | 200 | 198 | 986 |
| Total |  |  |  |  |  |  | 2681 |
| Primary Stem Width (PSW) | 1 | 146 | 156 | 155 | 156 | 177 | 790 |
|  | 2 | 166 | 180 | 175 | 199 | 186 | 906 |
|  | 3 | 200 | 190 | 199 | 200 | 197 | 986 |
| Total |  |  |  |  |  |  | 2682 |
| Flag Leaf Length (FLL) | 1 | 122 | 140 | 153 | 135 | 174 | 724 |
|  | 2 | 148 | 178 | 153 | 174 | 173 | 826 |
|  | 3 | 197 | 183 | 195 | 196 | 191 | 962 |
| Total |  |  |  |  |  |  | 2512 |
| Flag Leaf Width (FLW) | 1 | 128 | 145 | 153 | 147 | 172 | 745 |
|  | 2 | 145 | 175 | 159 | 180 | 173 | 832 |
|  | 3 | 198 | 184 | 196 | 200 | 195 | 973 |
| Total |  |  |  |  |  |  | 2550 |

**Supplemental Table S3.** Monthly weather data during field evaluations of cultivated Northern Wild Rice (cNWR; *Zizania palustris*) over three years, including average high and low temperatures for May through September of each year, as well as monthly average temperatures and average precipitation.

| Year | Month | Minimum<br>Temperature (C) | Maximum<br>Temperature (C) | Average<br>Temperature (C) | Average<br>Precipitation<br>(inches) |
| --- | --- | --- | --- | --- | --- |
| 2019 | May | 3.38 | 15.51 | 9.44 | 0.06 |
| 2019 | June | 10.40 | 23.22 | 16.81 | 0.20 |
| 2019 | July | 14.05 | 26.70 | 20.38 | 0.08 |
| 2019 | August | 9.27 | 23.34 | 16.30 | 0.08 |
| 2019 | September | 9.89 | 19.15 | 14.52 | 0.15 |
| 2020 | May | 5.39 | 17.59 | 11.49 | 0.03 |
| 2020 | June | 12.37 | 24.64 | 18.51 | 0.16 |
| 2020 | July | 15.32 | 27.55 | 21.43 | 0.17 |
| 2020 | August | 13.82 | 25.45 | 19.64 | 0.28 |
| 2020 | September | 7.33 | 18.08 | 12.70 | 0.06 |
| 2021 | May | 5.42 | 19.56 | 12.48 | 0.02 |
| 2021 | June | 13.32 | 27.49 | 20.40 | 0.04 |
| 2021 | July | 14.99 | 28.00 | 21.49 | 0.08 |
| 2021 | August | 14.79 | 26.39 | 20.59 | 0.06 |
| 2021 | September | 9.92 | 22.37 | 16.15 | 0.14 |

**Supplemental Table S4.** Yearly means and standard errors for basal stem width (BSW), flag leaf length (FLL), flag leaf width (FLW), plant height (PH), and primary stem width (PSW), measured over three years across five cultivated Northern Wild Rice (cNWR; *Zizania palustris*) populations. The “Average” column represents the across-population mean (± SE) for each trait in each year.

| Trait | Unit | Year | Barron | FY-C20 | Itasca-C12 | Itasca-C20 | K2 | Average |
| --- | --- | --- | --- | --- | --- | --- | --- | --- |
| **Basal Stem Width (BSW)** | mm | 1 | 14.93 ± 0.23 | 14.27 ± 0.23 | 14.37 ± 0.23 | 14.62 ± 0.23 | 12.42 ± 0.18 | 14.12 ± 0.22 |
| | | 2 | 10.01 ± 0.16 | 10.06 ± 0.15 | 10.84 ± 0.15 | 10.55 ± 0.14 | 12.52 ± 0.15 | 10.80 ± 0.15 |
| | | 3 | 14.11 ± 0.19 | 14.25 ± 0.19 | 14.76 ± 0.20 | 15.40 ± 0.21 | 14.30 ± 0.19 | 14.57 ± 0.20 |
| **Flag Leaf Length (FLL)** | cm | 1 | 33.97 ± 0.89 | 33.18 ± 0.84 | 37.20 ± 1.00 | 36.91 ± 0.90 | 31.69 ± 0.66 | 34.59 ± 0.87 |
| | | 2 | 30.91 ± 0.69 | 34.13 ± 0.76 | 33.49 ± 0.69 | 34.86 ± 0.70 | 38.26 ± 0.69 | 34.33 ± 0.71 |
| | | 3 | 39.65 ± 0.65 | 38.79 ± 0.80 | 37.62 ± 0.71 | 42.70 ± 0.79 | 37.20 ± 0.68 | 39.19 ± 0.73 |
| **Flag Leaf Width (FLW)** | cm | 1 | 2.51 ± 0.04 | 2.25 ± 0.04 | 2.45 ± 0.05 | 2.52 ± 0.05 | 2.33 ± 0.04 | 2.41 ± 0.04 |
| | | 2 | 1.92 ± 0.04 | 1.94 ± 0.04 | 2.04 ± 0.04 | 2.12 ± 0.04 | 2.58 ± 0.04 | 2.12 ± 0.04 |
| | | 3 | 2.87 ± 0.04 | 2.91 ± 0.05 | 2.45 ± 0.04 | 2.81 ± 0.04 | 2.56 ± 0.04 | 2.72 ± 0.04 |
| **Plant Height (PH)** | cm | 1 | 204.32 ± 2.06 | 190.28 ± 1.97 | 187.39 ± 2.19 | 186.57 ± 1.91 | 186.58 ± 1.87 | 191.00 ± 2.00 |
| | | 2 | 138.94 ± 1.55 | 140.42 ± 1.57 | 145.38 ± 1.54 | 153.63 ± 1.42 | 183.95 ± 1.36 | 152.46 ± 1.49 |
| | | 3 | 217.61 ± 1.43 | 218.14 ± 1.43 | 186.47 ± 1.31 | 201.14 ± 1.56 | 196.75 ± 1.30 | 204.02 ± 1.41 |
| **Primary Stem Width (PSW)** | mm | 1 | 6.63 ± 0.10 | 6.07 ± 0.09 | 6.48 ± 0.11 | 6.44 ± 0.10 | 5.91 ± 0.09 | 6.31 ± 0.10 |
| | | 2 | 5.31 ± 0.07 | 5.30 ± 0.08 | 5.59 ± 0.06 | 5.63 ± 0.08 | 6.34 ± 0.07 | 5.63 ± 0.07 |
| | | 3 | 7.39 ± 0.09 | 7.51 ± 0.09 | 6.86 ± 0.08 | 7.28 ± 0.07 | 7.27 ± 0.08 | 7.26 ± 0.08 |

**Supplemental Table S5.** Pearson correlation coefficients are shown for five morphological traits measured across all years combined. Asterisks (*) indicate statistically significant correlations (*P* < 0.05).

| Trait | PH | FLL | FLW | PSW |
| --- | --- | --- | --- | --- |
| PH |  |  |  |  |
| FLL | 0.22* |  |  |  |
| FLW | 0.50* | 0.67* |  |  |
| PSW | 0.55* | 0.62* | 0.75* |  |
| BSW | 0.59* | 0.31* | 0.51* | 0.60* |

**Supplemental Table S6.**

***See external excel file***

**Supplemental Figure S1.**
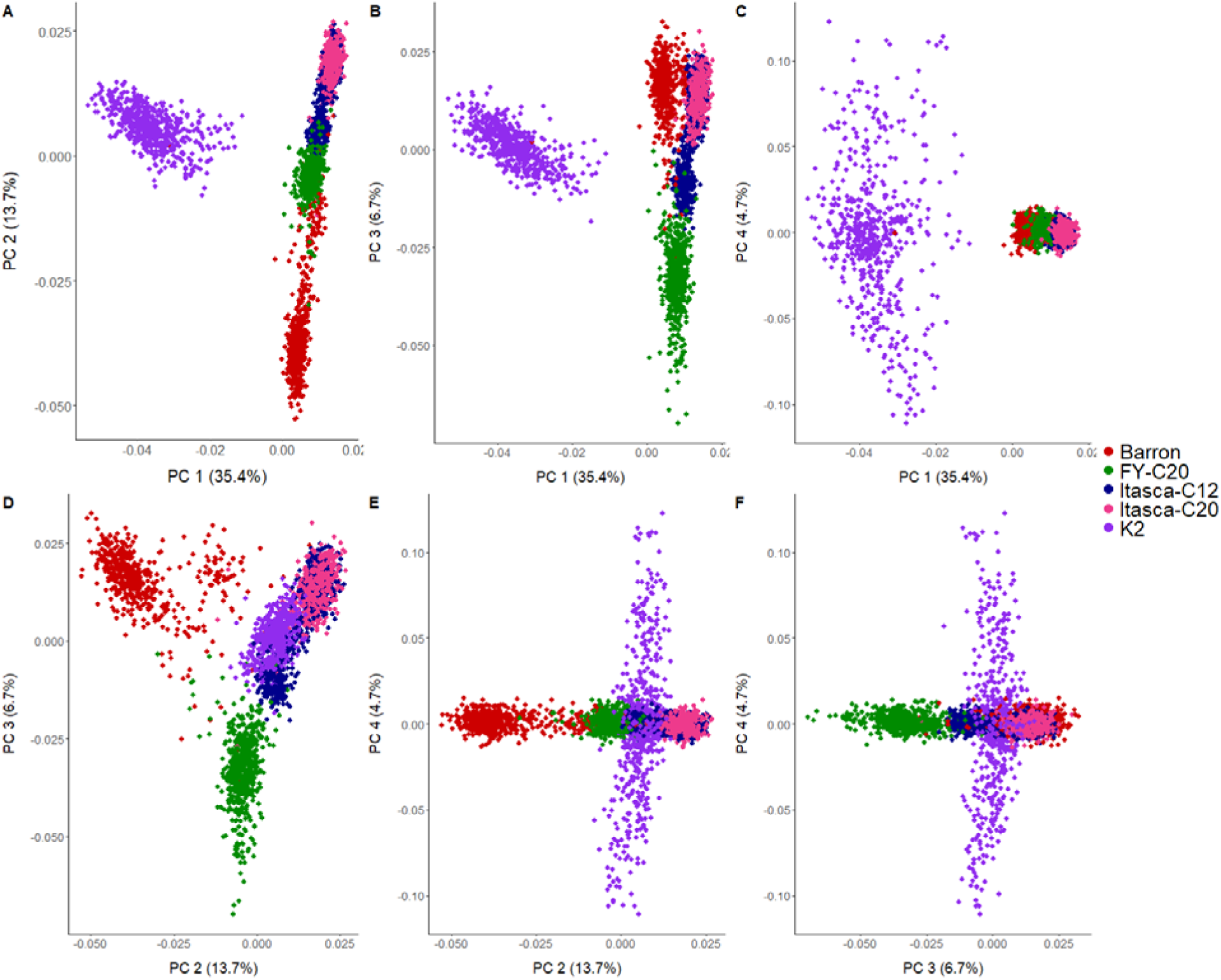
Principal component analysis (PCA) of cultivated Northern Wild Rice (*Zizania palustris* L.) populations across all years. Principal component analysis based on 73,363 filtered SNP markers from the combined multi-year dataset. Panels display pairwise combinations of the first four principal components: (A) PC1 × PC2, (B) PC1 × PC3, (C) PC1 × PC4, (D) PC2 × PC3, (E) PC2 × PC4, and (F) PC3 × PC4. Percent variance explained by each principal component is indicated on the axes. Each point represents an individual plant colored by population (Barron, FY-C20, Itasca-C12, Itasca-C20, and K2). PC1 consistently separates K2 from the remaining populations, while subsequent components resolve finer-scale differentiation among Barron and Itasca-derived populations.

**Supplemental Figure S2.**
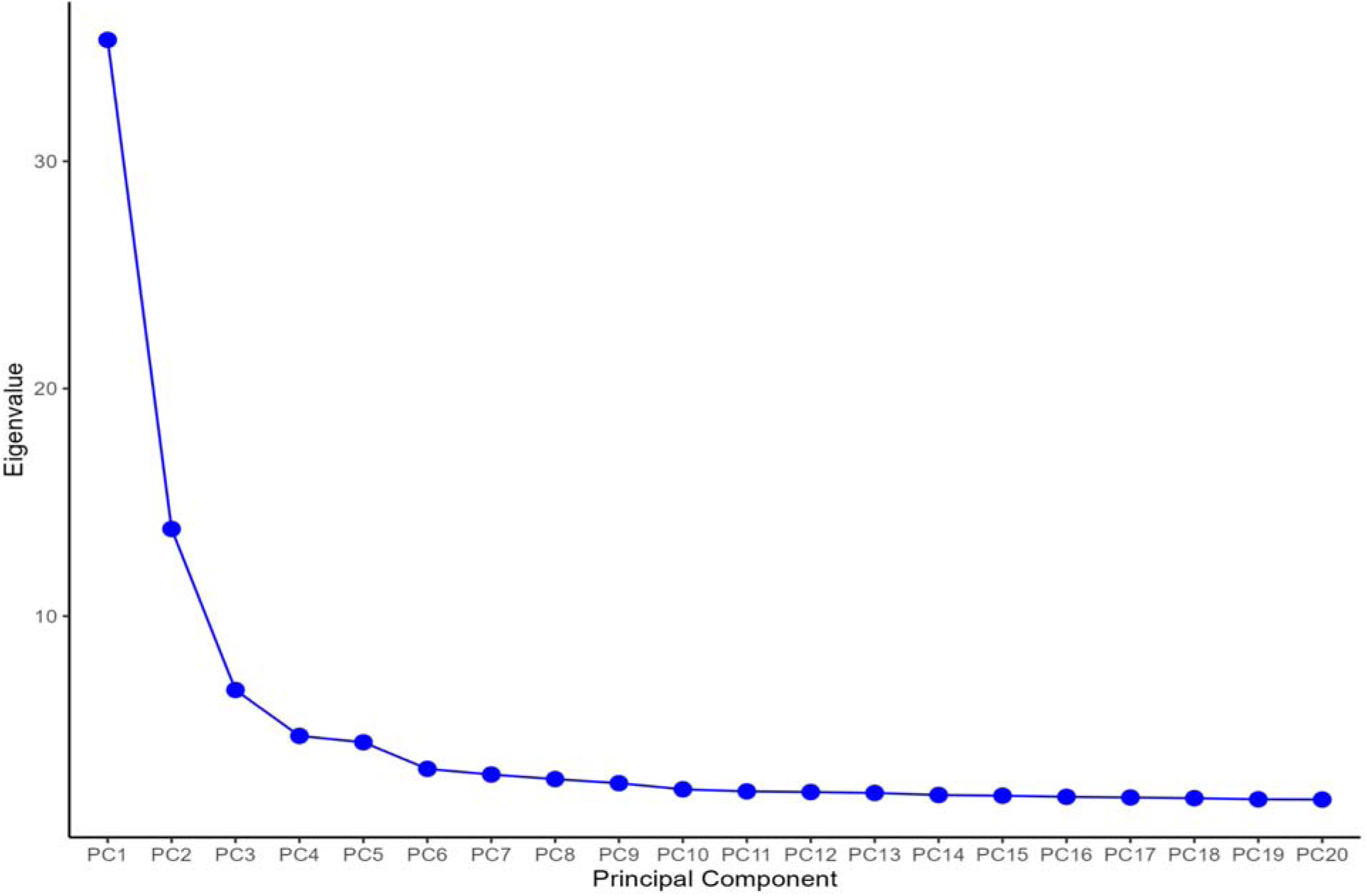
Scree plot showing the proportion of variance explained by each principal component for sp-GWAS data from 5 cultivated Northern Wild Rice (cNWR; *Zizania palustris*) populations across 3 years.

**Supplemental Figure S3.**
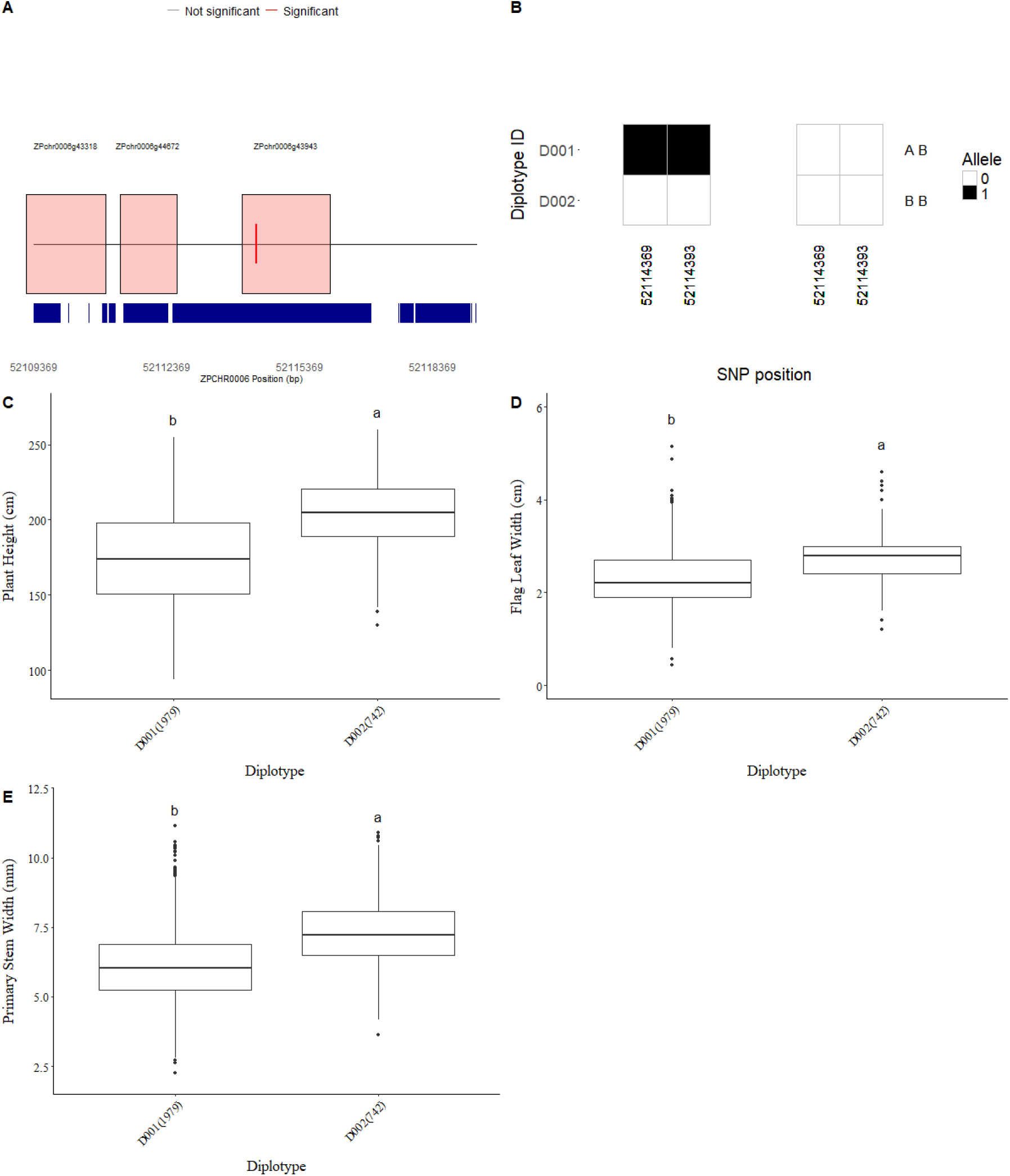
Diplotype structure and phenotypic effects at locus 6.16 in cultivated Northern Wild Rice (*Zizania palustris* L.), including (A) Local genomic region of locus 6.16 on chromosome 8, (B) Diplotype structure inferred from phased SNPs located within ±3 kb of the locus boundaries, haplotypes are labeled with capital letters on the far right (C-E) Phenotypic distributions of diplotype classes for (C) plant height (PH), (D) flag leaf width (FLW), and (E) primary stem width (PSW).

**Supplemental Figure S4.**
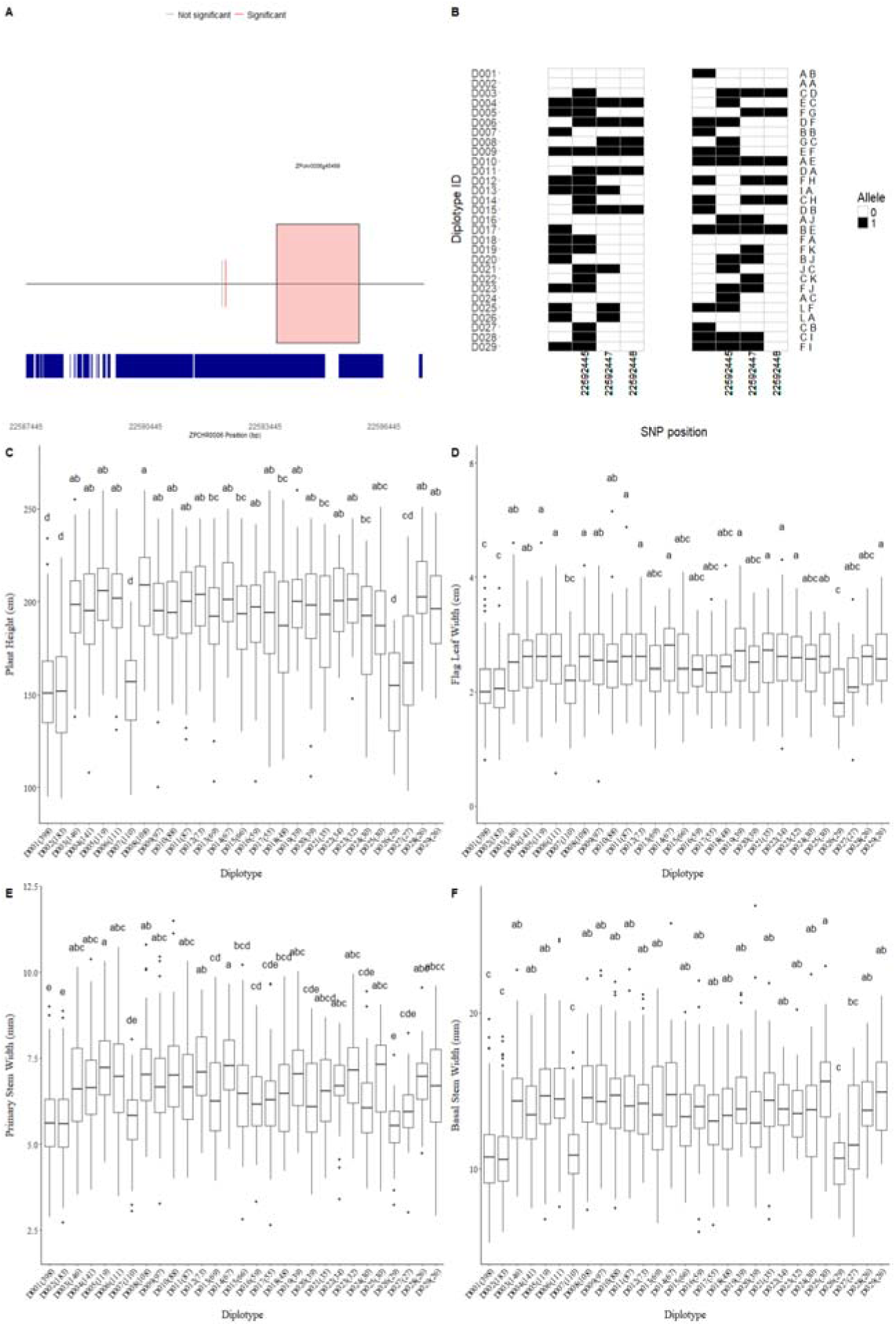
Diplotype structure and phenotypic effects at locus 6.3 in cultivated Northern Wild Rice (Zizania palustris L.), including (A) Local genomic region of locus on chromosome 6.3, (B) Diplotype structure inferred from phased SNPs located within ±3 kb of the locus boundaries, haplotype are labeled with capital letters on the far right of the graph, (C-F) Phenotypic distributions of diplotype classes for (C) plant height (PH), (D) flag leaf width (FLW), (E) primary stem width (PSW) and (F) basal stem width (BSW). Lower case letters above denote statistically significant differences among diplotype based on Tukey’s honestly significant difference (HSD) test (α = 0.05)

**Supplemental Figure S5.**
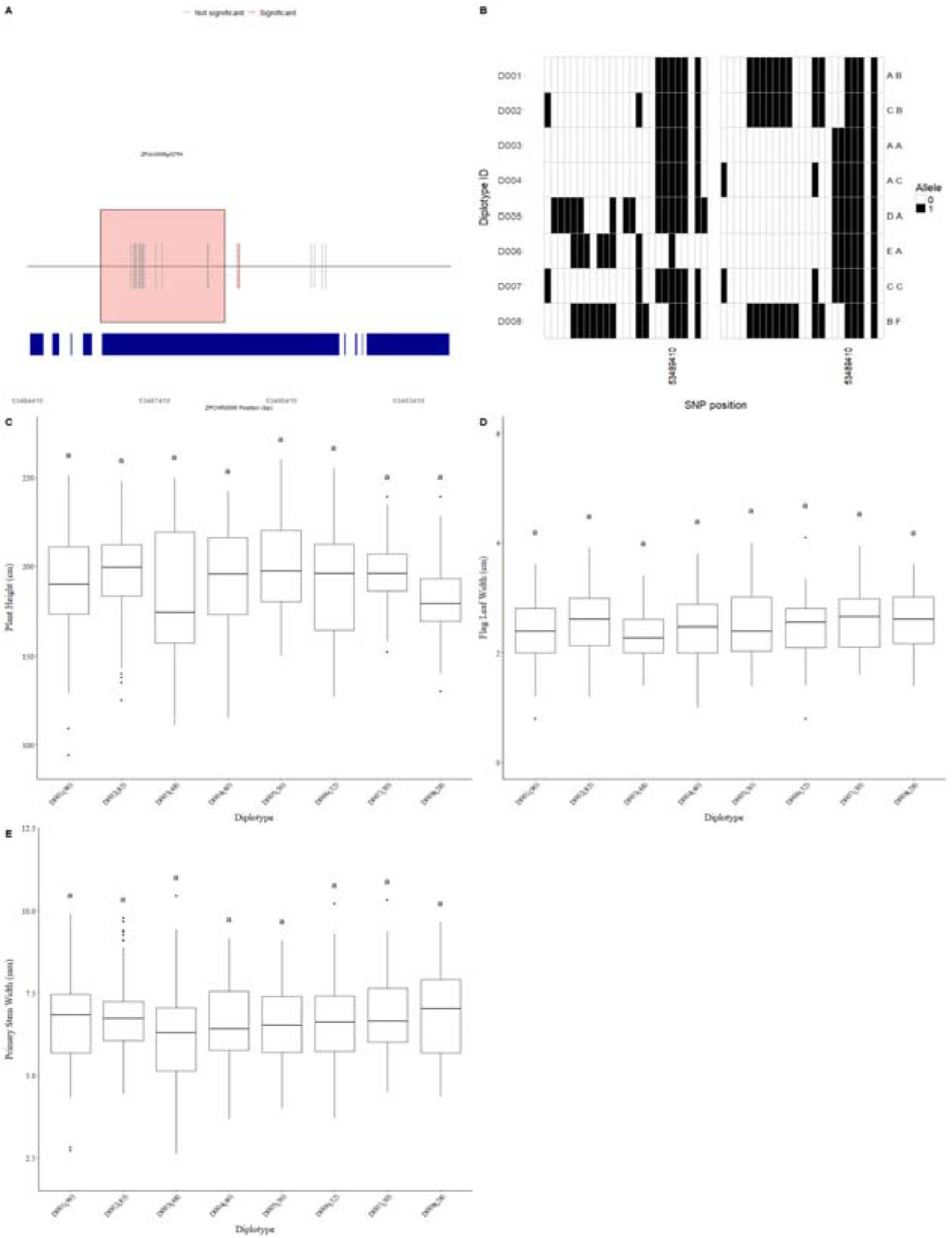
Diplotype structure and phenotypic effects at locus 6.18 in cultivated Northern Wild Rice (Zizania palustris L.), including (A) Local genomic region of locus 6.18 on chromosome 8, (B) Diplotype structure inferred from phased SNPs located within ±3 kb of the locus boundaries, haplotype are labeled with capital letters on the far right of the graph, (C-E) Phenotypic distributions of diplotype classes for (C) plant height (PH), (D) flag leaf width (FLW), and (E) primary stem width (PSW). Lower case letters above denote statistically significant differences among diplotypes based on Tukey’s honestly significant difference (HSD) test (α = 0.05)

**Supplemental Figure S6.**
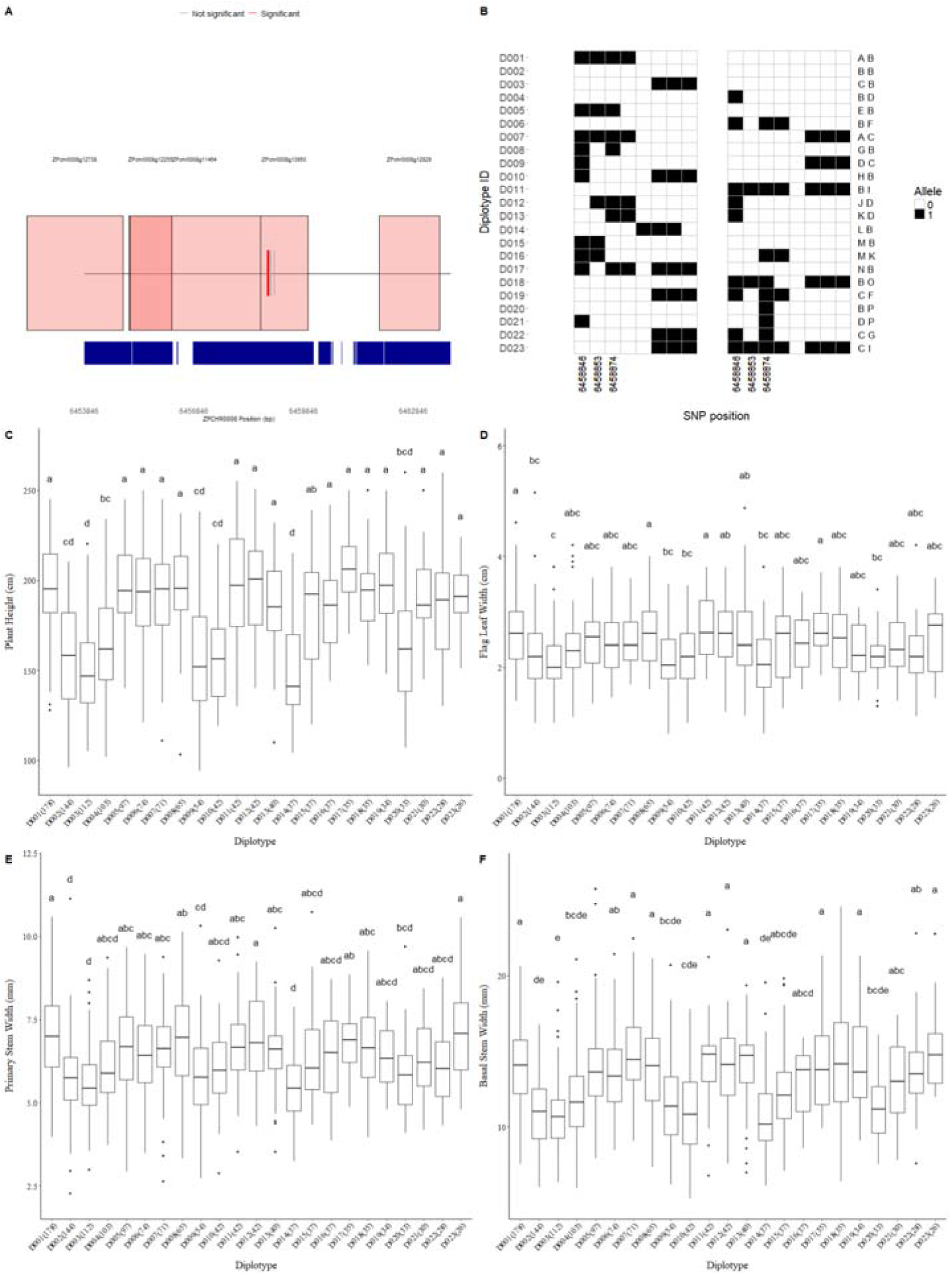
Diplotype structure and phenotypic effects at locus 8.3 in cultivated Northern Wild Rice (Zizania palustris L.), including (A) Local genomic region of locus 8.3 on chromosome 8, (B) Diplotype structure inferred from phased SNPs located within ±3 kb of the locus boundaries, haplotype are labeled with capital letters on the far right of the graph, (C-E) Phenotypic distributions of diplotype classes for (C) plant height (PH), (D) flag leaf width (FLW), (E) primary stem width (PSW) and (F) basal stem width (BSW). Lower case letters above denote statistically significant differences among diplotype based on Tukey’s honestly significant difference (HSD) test (α = 0.05)

